# Morphometrics and phylogenomics of coca (*Erythroxylum* spp.) illuminate its reticulate evolution, with implications for taxonomy

**DOI:** 10.1101/2023.11.28.569066

**Authors:** Natalia A. S. Przelomska, Rudy A. Diaz, Fabio Andrés Ávila, Gustavo A. Ballen, Rocío Cortés-B, Logan Kistler, Daniel H. Chitwood, Martha Charitonidou, Susanne S. Renner, Oscar A. Pérez-Escobar, Alexandre Antonelli

## Abstract

South American coca (*Erythroxylum coca* and *E. novogranatense*) has been a key-stone crop for many Andean and Amazonian communities for at least 8,000 years. However, over the last half century, global demand for cocaine has placed this plant in the centre of armed conflict, deforestation, and explosive growth of illegal economies. While national and international agencies progress from a ‘war on drugs’ policy model towards locally appropriate, data-informed strategies to tackle coca plantations, monitoring their expansion and composition remains essential. The principal means to identify coca plants is leaf morphology, yet the extent to which it is reflected in taxonomy is uncertain. Here, we analyse the consistency of the current naming system of coca and its four closest wild relatives (the ‘coca clade’), using morphometrics, phylogenomics, and population genomics. We include the name-bearing type specimens of coca’s closest wild relatives *E. gracilipes* and *E. cataractarum*. Morphometrics of 342 digitized herbarium specimens show that leaf shape and size fail to reliably discriminate between species and varieties. However, the rounder and more obovate leaves of certain coca varieties could be associated with domestication syndrome of this crop. Our phylogenomic data indicate gene flow involving monophyletic clades of *E. gracilipes* and the *E. coca* clade. These results further clarify the evolution of coca and support a taxonomic framework wherein *E. gracilipes* is retained as a single species. Our findings have implications for the development of cost-effective genotyping methods to effectively discriminate varieties of cultural significance from high-yielding cultivars fuelling the lucrative cocaine market.

## Introduction

The expansion of humans into South America some 15,000 to 13,500 years ago (Rothhammer & Dillehay, 2009) was accompanied by the adoption, and sometimes domestication of native plants into human agroecosystems. One of the earliest known and most widely cultivated crops was coca (Rury & Plowman, 1984) (*Erythroxylum coca* and *E. novogranatense*; Schulz, 1907; Machado 1970; Plowman, 1979), whose history of use traces back at least 8,000 years (Dillehay et al., 2010). Coca was predominantly involved in the cultural evolution of societies owing to its medicinal, stimulatory and cultural properties (Plowman, 1984; Schultes, 1979), and many Andean and Amazonian communities today remain reliant on coca (Azevedo, 2021; Cristancho & Vining, 2004; Naranjo, 1979; Vergara et al., 2022). Despite many ethnobotanical and physiological studies, the taxonomic boundaries between cultivated varieties and their wild relatives in the genus *Erythroxylum* (family Erythroxylaceae) are poorly defined. The implications of an inadequate classification system of coca are pertinent to illegal plantations. Blanket eradication policies, deforestation and armed conflict stemming from the decades-long ‘war on drugs’ have become associated with such plantations (Negret et al., 2019; Rincón-Ruiz & Kallis, 2013) and have also played a part in threatening the Indigenous biocultural diversity of coca by uprooting natural plantations. Such misdirected activity could be curtailed if monitoring systems focussing on cocaine production plantations involved a workable plant identification procedure before considering eradication.

The genus *Erythroxylum* consists of over 270 species, of which about three quarters are native to the American tropics (Jara-Muñoz et al., 2022; Plowman & Hensold, 2004).

There are four varieties of cultivated cocas – two each in the species *Erythroxylum coca* and *E. novogranatense*, which have largely allopatric distributions in northwestern South America (Bohm et al., 1982; our Fig. 1). The more widely cultivated species is *E. coca* (Huánuco coca). Its variety *coca* is native to wet montane forests of the eastern Andean slopes of Peru and Bolivia and the other variety *ipadu* Plowman (Amazonian coca) grows across the lowland Amazon basin. The less widely cultivated *E. novogranatense* (D. Morris) Hieron. has historically been grown in the dry valleys of the Cordilleras and the Sierra Nevada de Santa Marta (Bohm et al., 1982), but also recently found in the Pacific region (Chocó and Cauca) (UNODC 2016). Its variety Trujillo coca (var. *truxillense* (Rusby) Plowman) is cultivated in arid regions of north-western Peru for traditional use and is a flavouring and stimulant additive to the soft drink *Coca Cola* (Plowman, 1986; Gootenberg, 2003).

**Figure 1.**
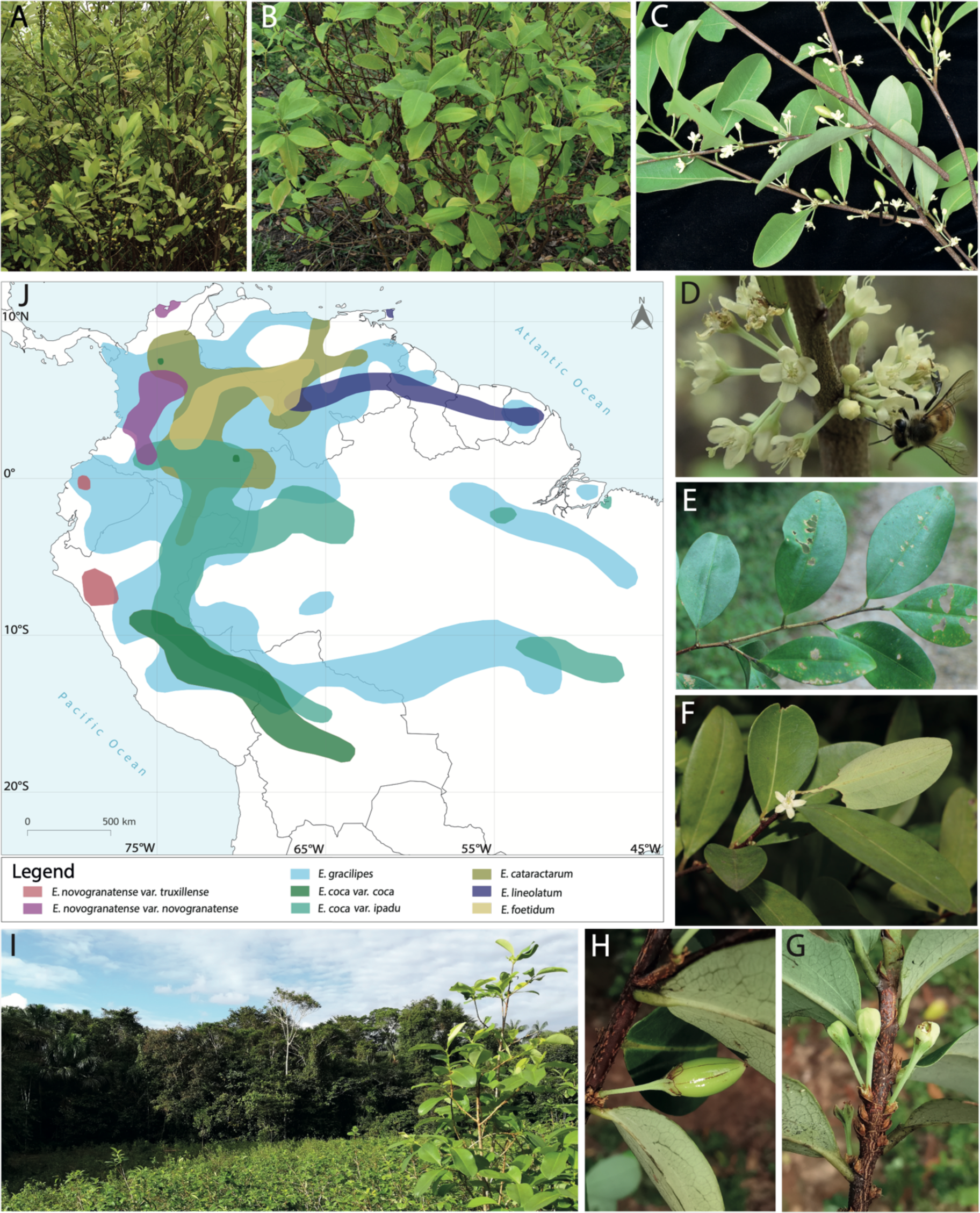
Morphology and geographical distribution of taxa in the coca clade. (A) Bush of *E. novogranatense.* **(B)** Bush of *E. coca* var. *coca.* **(C)** Leaves, flowers, immature flowers and immature fruit of *E. coca* var. *ipadu.* **(D)** Flowers of *E. coca* var. *ipadu* with potential pollinator. **(E)** Leaves of *E. gracilipes.* **(F)** Leaves of *E. cataractarum.* **(G)** Immature flowers of *E. cataractarum.* **(H)** Immature fruit of *E. cataractarum.* **(I**) Illegal coca plantation established in the Amazonian forests of Putumayo department, Colombia. **(J)** geographical distributions of cultivated and wild relative *Erythroxylum* taxa. Photos: Rocío del Pilar Cortés Ballén (A,B,C,D), William Ariza (E,F), José Aguilar Cano (G,H,I).

Attribution of species status to the two broad types of cultivated coca was undoubtedly influenced by their ethnobotanical significance, and differences between them were discerned through morphology, chemotaxonomy, and mating systems (Bohm et al., 1982; Plowman 1986; Plowman & Rivier, 1983; Rury, 1981). A better understanding of the relatedness of the lineages of cultivated crops can now be gained with molecular markers (Viruel et al., 2021), and population-level genomic work (White et al., 2021) has suggested that cultivated coca is polyphyletic, with wild *E. gracilipes* inferred as the wild progenitor of both species.

As a leaf crop, selective pressures may have affected coca leaf shape and size to the point of these becoming taxonomically informative; leaf morphology can be part of the domestication syndrome (Arias et al., 2021; Galindo Bonilla & Fernández-Alonso, 2010). Indeed, leaf characteristics have been crucial in the formal taxonomy of coca (Plowman, 1979; Plowman, 1982), an approach still used today for coca identification in monitoring surveys (eg. UNODC, 2012). The leaves of cultivated coca are thought to be distinguishable from leaves of closely related, and often sympatric wild *Erythroxylum* species by being smaller, rounder and softer (Rury, 1981; White et al., 2021). However, intergrading variation across the two cultivated coca species and their wild relatives is pervasive (Rury & Plowman, 1984), and phenotypic plasticity additionally renders leaf morphology-based identification of individual coca leaves problematic (Rury, 1981; Rury & Plowman, 1984). Rigorous statistical comparisons so far are lacking.

Here, we investigate genetic relationships in the coca clade, assess their matching with taxon boundaries and examine the discriminatory power of leaf morphology in identifying species and varieties using a large sample of herbarium specimens. We employ Gaussian Mixture Models (GMMs) to infer probabilistic morphometric clusters and assess their overlap with the currently accepted taxa. We then infer population-level nuclear and plastid phylogenies for the coca clade through a hybrid approach of genome skimming herbarium specimens and by mining published *Erythroxylum* target capture genomic datasets. To test for gene flow among the taxa, we apply phylogenetic network analysis. Finally, we apply population genomic tests and molecular-clock models to circumscribe population groups of coca and estimate lineage divergence times.

## Results

### Leaf size is insufficient for identifying the cultivated species of coca

We find that leaf size metrics have limited power to discriminate between wild relatives and cultivated cocas. In our principal component analysis (PCA) on linear metrics, the taxa do not appear as distinct clusters. Nevertheless, PC1 does segregate most *E. gracilipes* individuals from the remaining taxa (Fig. 2A) due to the greater leaf area (loadings for area, width and length: 0.590, 0.574 and 0.572). Leaf area is likewise significantly greater in *E. foetidum* compared to the remaining taxa (t-test, *P*<2.2e-4) (Figs. 2A, S1, S4). PC2 can be attributed to the leaf length-to-width ratio (loadings: -0.73 and 0.68 respectively, loading of area: 0.04). We noted taxonomic signal for length-to-width ratio, with diverging slopes for different taxonomic groupings on a LOESS plot (Fig. S5). *E. novogranatense* var. *truxillense* has longer, narrower leaves than those observed in *E. n.* var. *novogranatense*. *E. coca* in turn has slightly larger leaves (Figs. 2,S6). However, the varietal groups of *E. novogranatense* and *E. coca*, collectively with *E. cataractarum*, exhibit a large proportion of overlap in leaf morphospace (Figs. 2A,B), precluding statistically supported morphogroup delimitation. The *E. cataractarum* type specimen appears near the extreme end of its morphospace (Fig. 2A), and closer to the centroid of *E. n.* var. *truxillense*’s distribution in this morphospace.

**Figure 2.**
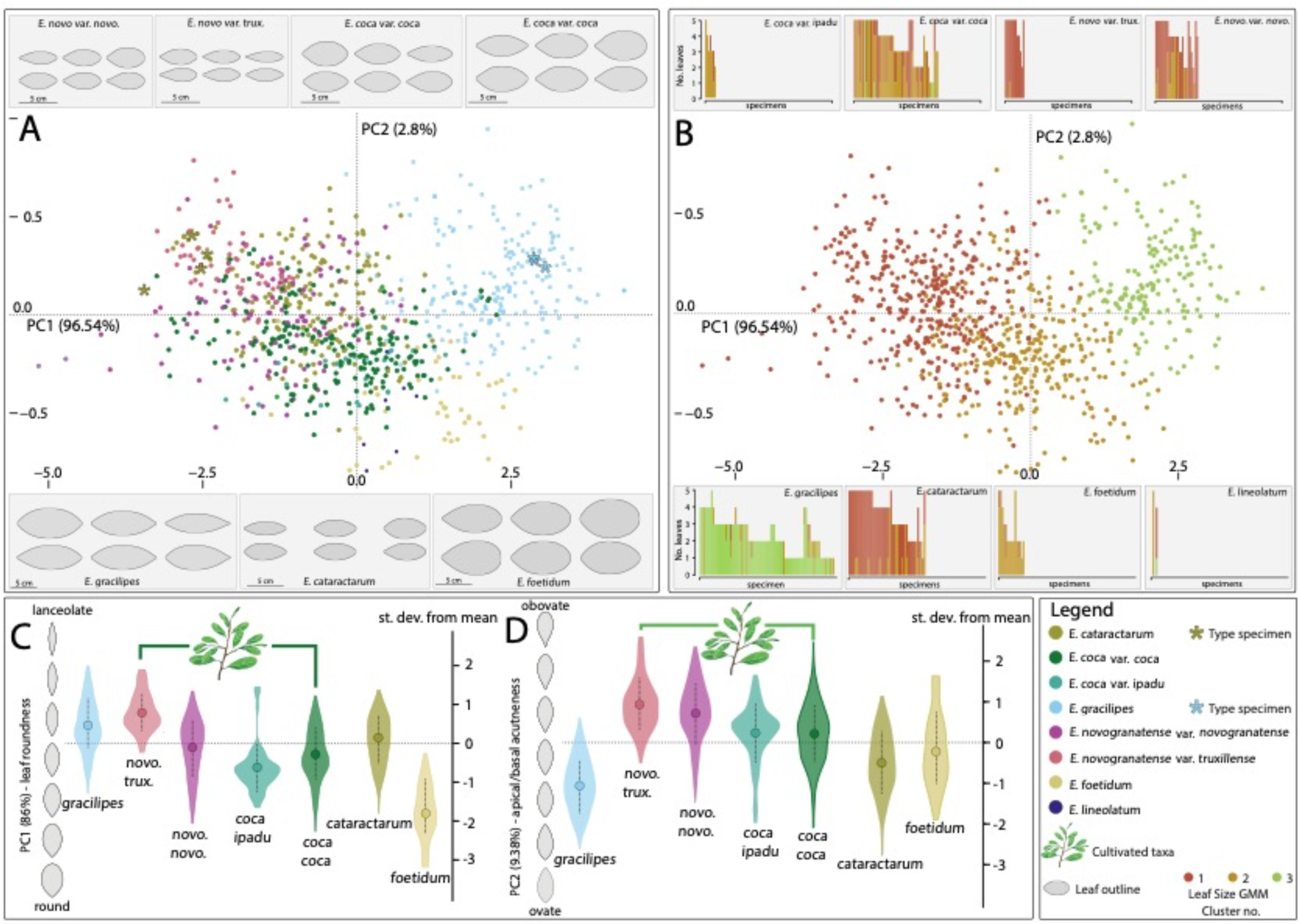
**Summary of leaf morphometric analyses, encompassing leaf size and leaf shape**. **(A)** Principal component analysis (PCA) of morphospace based on linear metrics, coloured by verified pre-defined taxonomic identifications. Insets are eigen-leaves computed for each taxonomic subset of the data, reconstructed using the reverse Fourier transform for the mean, and +/1.5 standard deviations from the mean, on the first two PCs of shape variation within a species. **(B)** PCA of morphospace based on linear metrics, coloured by assignment resulting from the GMM clustering model. Insets are bar charts representing the number of leaves sampled, and their individual cluster assignments, grouped by pre-defined taxonomic identifications. **(C)** Plots of the distribution of values along the first PC of elliptical Fourier analysis (EFA) PCA representing leaf roundness, grouped by pre-defined taxonomic identification **(D)** Plots of the distribution of values along the second PC of elliptical Fourier analysis (EFA), representing acuteness at the base or apex, grouped by pre-defined taxonomic identification. *E. lineolatum* was omitted from the EFA shape analyses due to limited sample size.

The GMMs with the highest level of statistical support model three clusters for allometric data (Fig. 2B), with consistent support even after downsampling to 50 leaves per taxon and inclusion of samples grown either outside of their native South American range or in cultivation (Figs. S1-3). Most of the correspondence between clustering patterns that was detected was likely driven by cluster 3, which exhibits a good fit to the single taxonomic group of *E. gracilipes.* Group 2 was a good fit for *E. foetidum* and *E. coca* var. *ipadu* and most common in *E. coca* var. *coca*, whereas Group 3 is a good fit for *E. novogranatense* var. *truxillense*, and the dominant group for *E. n.* var. *novogranatense* and *E. cataractarum*. Importantly, these clustering methods based on linear metrics fail to distinguish the three wild relative species consistently and confidently from cultivated cocas. This linear metric-based clustering was discordant with the taxonomic groupings overall (Rand index, full dataset = 0.69, Rand index, cultivated cocas only = 0.632; Table S4).

### Leaf shape morphometrics illuminate traits that define cultivated species

Using a reverse Fourier transform, we directly visualise outlines of leaf shapes using the statistical framework of the PCA into which our raw leaf outline data were input (Figs. 2A; S1E,F; S2E,F; S3E,F), revealing the most defining leaf shape traits (Fig. 2A,C,D,S1-3,S9). PC1 describes leaf elongation from obovate to lanceolate, or ‘roundness’ of the leaves. *E. foetidum* has the roundest leaves, followed by *E. coca* var. *ipadu* and *E. coca* var. *coca*, whereas *E. novogranatense* var. *truxillense* has the most lanceolate leaves (Fig. 2C). PC2 describes obovate versus ovate shape of the leaves, or ‘acuteness at the base or apex’ (Fig. 2D), where wild taxa (*E. gracilipes*, *E. cataractarum* and *E. foetidum*) have more ovate leaves than the cultivated species (t-test, *P*<2.2e-16). This trait is most pronounced in *E. gracilipes*. The cultivated varieties are better characterised by obovate leaf shape, and especially *E. novogranatense* var. *truxillense*. As with the linear metric data, our GMM of highest statistical support clusters the data into three groups (consistent after downsampling), albeit with differences in group composition. Aspects of congruence between shape and size GMMs clusters are as follows. One of the modelled groups (Group 3) is assigned almost exclusively to *E. gracilipes* individuals. The principal group to which *E. foetidum* individuals is assigned (Group 1) is in turn distinct from this. Group 2 is prevalent, dominating in all cultivated taxa and *E. cataractarum*, but also common in *E. foetidum* and *E. gracilipes* (Fig. S8). The PCoA based on EFA data likewise shows how all cultivated varieties and *E. cataractarum* form a close cluster (Fig. S9). Overall, there is less uniformity of taxa when grouped using shape data compared to when grouped based on linear metric data, and an even greater mismatch of GMM-inferred clusters to the taxonomy (Rand index, full dataset = 0.555, Rand index, cultivated cocas only = 0.425; Table S5).

### E. gracilipes gene flow is pervasive and supported by hybrid edges

The uniparental plastid phylogeny and the 326-gene nuclear phylogeny both underscore the complex genetic structure of *E. gracilipes* (Figs. 3A-D), and each genomic source places *E. foetidum* (and sample ‘Spruce 3725’) as sister to the coca clade. The remaining portions of the respective phylogenies exhibit incongruence, especially in the placement of the 10 *E. gracilipes* specimens, including the type specimen Spruce 3068, which clusters with *E. novogranatense* in the plastid tree and with *E gracilipes* + *E. coca* in the nuclear tree.

**Figure 3.**
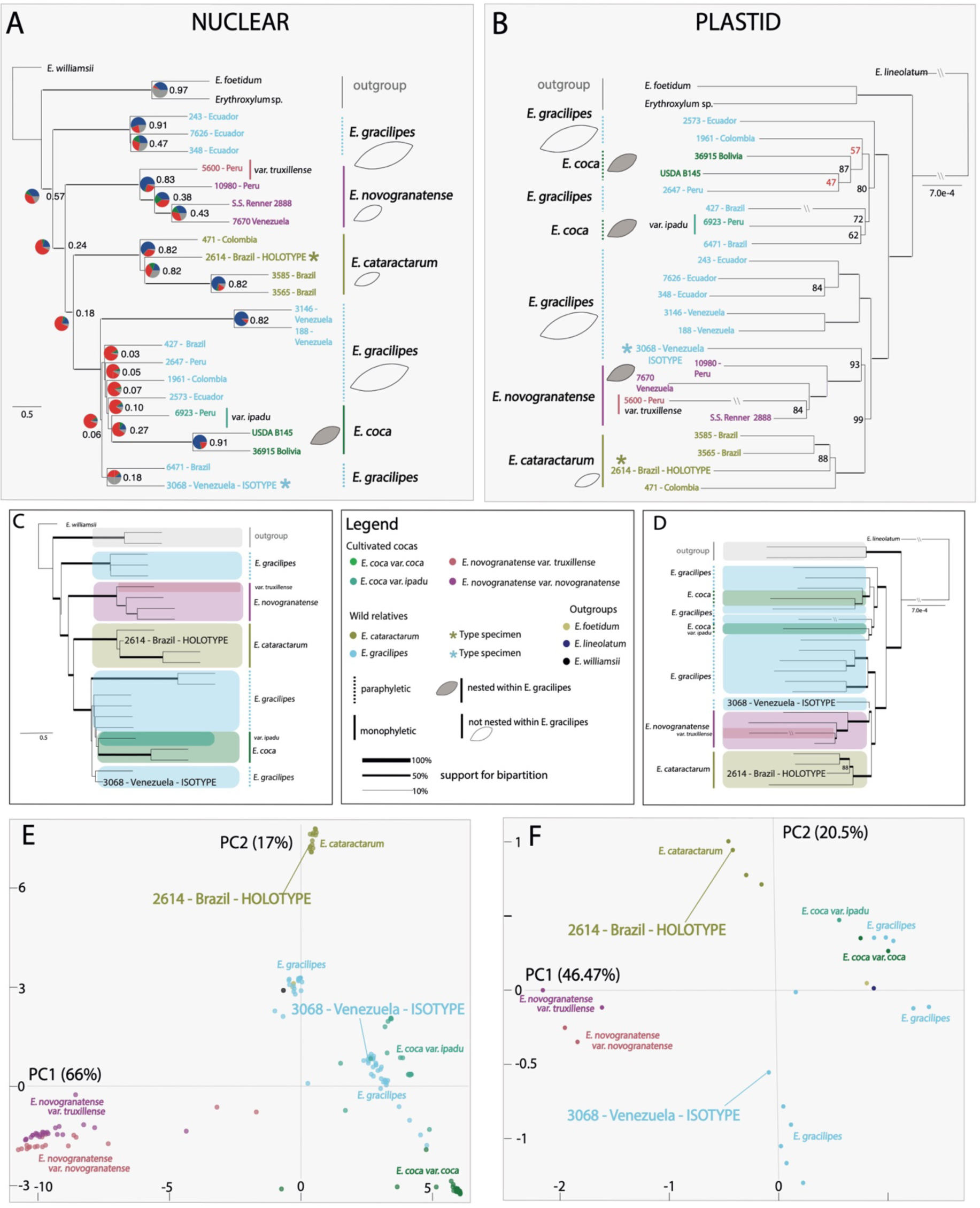
**Comparative phylogenomic and population genomic analyses of coca and wild relatives**. **(A)** ASTRAL summary species tree topology derived from 326 maximum likelihood (ML) phylogenies built from nuclear genes from 25 samples of *Erythroxylum*, with percent quartet support indicated as branch thickness and gene concordance factors presented. Pie charts at nodes indicate the proportion of gene trees that support (blue), or conflict with (green, red) the species tree. Here, the green proportion of the pie chart indicates the proportion of gene trees recovering the second most frequent alternative bipartition to the species tree, and red – any other alternative conflicting bipartition. Grey represents non-informative gene trees. **(B)** ML tree reconstructed from plastomes sequenced in 25 samples of *Erythroxylum,* with bootstrap support after 500 iterations indicated and bootstrap support represented as branch thickness. **(C)** ASTRAL summary species tree topology derived from 326 maximum likelihood (ML) phylogenies, with clades highlighted and colour-coded according to taxon and type specimens indicated. **(D)** ML tree reconstructed from plastomes, with clades highlighted and colour-coded according to taxon and type specimens indicated. **(E)** Nuclear genomic PCA based on 13,643 genotype likelihoods (GLs) computed from 25 sequenced and 148 data-mined accessions of *Erythroxylum* samples. Sequenced type specimens for *E. cataractarum* and *E. gracilipes* highlighted. **(F)** Plastid PCA based on 4,664 GLs computed from sequenced and datamined accessions of 25 *Erythroxylum* samples. Sequenced type specimens for *E. cataractarum* and *E. gracilipes* highlighted in E and F.

Regarding *E. coca*, the plastid tree infers closely related but phylogenetically distinct *coca* and *ipadu* varieties within a poorly resolved *E. gracilipes* + *E. coca* clade. In the nuclear dataset, the multispecies coalescent ASTRAL species tree indicates that gene tree incongruence is substantial (normalised quartet score: 0.657). For any given bipartition within this *E. gracilipes* + *E. coca* clade, there is a mean total of 12 gene trees supporting the bipartition with strong support and 94 gene trees conflicting with strong support (Fig. 3A) (mean gene concordance factor (gCF)=0.11, mean internode certainty (IC)=0.08, Figs. S12a-d). Within this, a monophyletic clade of *ipadu* and *coca* varieties has moderate support (32 supporting trees, 78 conflicting trees, gCF=0.27, IC=0.28), but includes a highly supported sister status of the two *coca* samples (gCF=0.91, IC=0.86). In contrast, *E. cataractarum* and *E. novogranatense* are well-supported, monophyletic groups, with 100% and 93% BS support in the plastid phylogeny, and characterised by high gene tree support values in the ASTRAL tree (*E. cataractarum* gCF=0.86, IC=0.82; *E. novogranatense* gCF=0.87, IC=0.83).

The phylogenetic network analysis, conducted to gain a broader picture of the evolutionary history of the clade, provides support to the hypothesis of gene flow involving *E. gracilipes.* The best network suggests two reticulations (Fig. S14). The first is highly supported (BS value = 98) and originates in the same clade as the early-diverging, monophyletic Ecuadorian *E. gracilipes* clade, with the *E. gracilipes* + *E. coca* clade as the recipient (inheritance proportion = 0.704). The second hybridization edge detected sits within the *E. gracilipes* + *E. coca* clade, and likewise involves a monophyletic *E. gracilipes* outgroup supplying genetic material to an ingroup including the cultivated lineages (inheritance proportion = 0.827). An alternative recipient position with lower support was recovered for this second hybridisation. The main source of this uncertainty likely lies with the recipient branch, on the basis that the donor branch has a higher combined support (76). However, since the orientation of gene flow relies on support of the minor edge in the hybridisation event, and is concordant here, we consider gene flow direction to be highly supported.

### Bioculturally important varieties differ in degree of intraspecific genetic resolution

Our PCA analyses based on nuclear genotype likelihoods (GLs) reveal extensive genetic diversity within *E. gracilipes*. The most substantial proportion of this diversity manifests as a cluster containing the K isotype of *E. gracilipes* (type: Spruce 3068), corresponding to ‘gracilipes1’ *sensu* White et al. (2021). This group, along with a secondary *E. gracilipes* and a single *E. cataractarum* cluster, are observed in both the merged dataset (Fig. 3E) and in our wild relatives-focussed sampling from Kew (K) specimens only (Fig. S15). The merged dataset PCA (Fig. 3E) exhibits the largest proportion of the variation as being driven by *E. novogranatense* and *E. coca*. Within *E. novogranatense,* its two varieties do form separate clusters, but with more subtle differentiation. In comparison, the varieties of *E. coca*: var. *coca* and var. *ipadu* are distinct. Unsupervised clustering predicted highest values of Δ*K* for scenarios of genetic structuring where *E. novogranatense* (both varieties), *E. coca* var*. coca*, *E. gracilipes* (containing *E. coca var. ipadu*) and *E. cataractarum* resolve as 3 or 4 broadly separate genetic entities (Figs. 4A,5B, S16, S17). The isotype-defined *E. gracilipes* group becomes distinct at *K*=5 (Fig. 4B) and *E. coca* var. *ipadu* at *K*=6 (Fig. S15). Only under a model of 8 hypothetical populations does *E. novogranatense* var. *novogranatense* exhibit genetic differentiation from *E. novogranatense* var. *truxillense* (Fig. S16).

**Figure 4.**
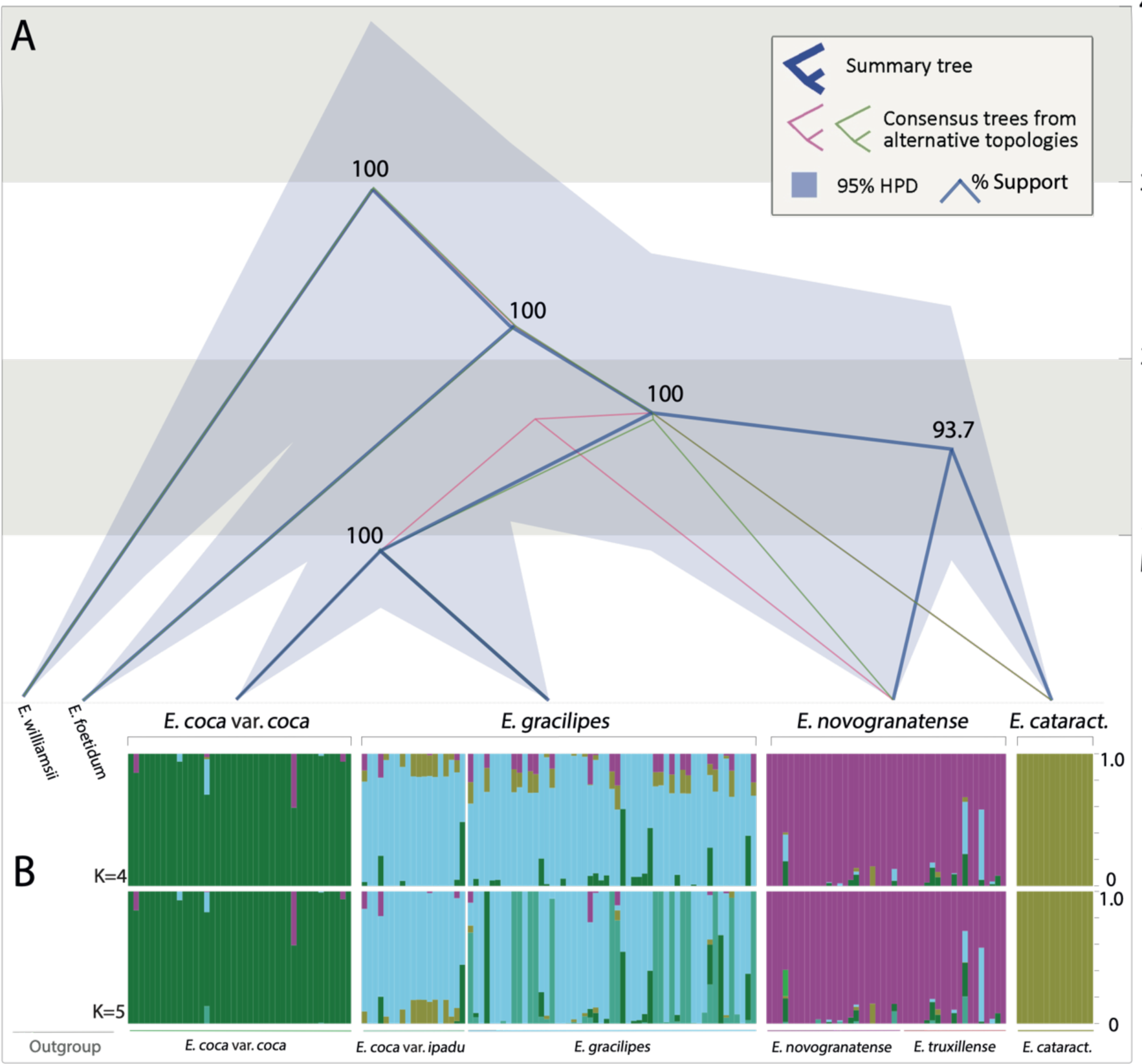
Absolute times of divergence of lineages in the coca clade based on populations. (A) Chronogram representing dates of divergence within the clade currently including *E. gracilipes*, *E. cataractarum*, *E. coca* and *E. novogranatense*. *E. foetidum* and *E. williamsii* serve as successive outgroups. Consensus topology shown in dark blue. Alternative topologies in pink and green. 95% Highest Posterior Density Intervals of absolute ages indicated as shaded area and % support provided at each node. **(B)** NGSadmix population structure inference showing per-individual ancestry of sequenced and data-mined accessions of 173 *Erythroxylum* specimens at highly supported values of *K*=4 and *K*=5. Full results of per-individual ancestry inference for varying values of *K* provided in supplementary materials.

### Lineage divergence of cultivated cocas long precedes the peopling of South America

We inferred a time-calibrated phylogeny using a Bayesian multispecies coalescent (MSC) to estimate of ages of divergence within this clade. In our population-focussed approach, we combined a species tree relaxed molecular clock model with information on population membership determined using NGSadmix. We reliably reconstructed a species tree for the cultivated cocas and inferred absolute ages of their divergence from close wild living relatives in the presence of topological discordance (Ogilvie et al., 2017). The results imply that the coca clade diversified 2.2 Ma (+/- 1 Ma, PP = 1.0) (Fig. 4A). The cultivated cocas and their wild living relatives shared a common ancestor ∼1.6 Ma ago (+/- 700 Kyrs, PP = 1.0). The divergence of the cultivated *E. novogranatense* and *E. coca* var. *coca* lineages from *E. cataractarum* and *E. gracilipes* (1400 - 800 Kyrs, +/- 800 - 400 Kyrs, 0.93 and 1.0 PP, respectively) precedes the peopling of the American tropics (∼15,5Kya; Dillehay et al., 2008; Prates et al., 2020; Rothhammer & Dillehay, 2009) by hundreds of millennia.

## Discussion

Establishing a robust taxonomic framework is vital for research into plants that are medicinally, nutritionally or culturally valuable to humans. This is because plant classification at this scale involves elucidating the plants’ evolutionary trajectories within a wider pool of genetic diversity comprising crop wild relatives, hybrids and semi-domesticated forms (Pellicer et al., 2018; Pérez-Escobar et al., 2021; Simon et al., 2022). A point in case is coca, a plant so highly prized for its tropane alkaloids that it has become a lynchpin of socioeconomic chaos – the naming system of coca is directly linked to legal ramifications around its cultivation and trafficking.

Our new plastid tree for the coca clade corroborates the hypothesis of White et al. (2021) proposing *E. gracilipes* as a clade within which *E. cataractarum*, *E. novogranatense* and *E. coca* are nested. Our population genomic and phylogenomic analyses which build upon this study illustrate extensive genetic structure characterizing *E. gracilipes*, a shrub proposed as the wild progenitor of all described varieties of cultivated coca (Macbride 1949; White et al 2021). It is unsurprising that a plant species with a broad geographic distribution should show non-discrete clustering (Dodsworth et al., 2021) and *E. gracilipes* is indeed distributed widely – in wet biomes across tropical South America (White et al., 2019) (Fig. 1J). Counteracting this structure is prevalent gene flow between lineages currently considered as *E. gracilipes*, involving both early diverging clades and the admixed *E. gracilipes + E. coca* clade. This contrasts to the findings of White et al (2021) who conducted maximum likelihood Treemix analysis (Pritchard & Pickrell, 2012) to conclude that none of the three most significant waves of gene flow in the coca clade involved the poorly resolved *E. gracilipes + E. coca* clade, but instead featured *E. cataractarum* as donor or recipient.

Our largest *E. gracilipes* cluster defined by the Spruce 3068 isotype, was previously identified as ‘gracilipes1’ (White et al. 2021), who proposed to retain it as a genetically constricted species of *E. gracilipes*, while reclassifying other pockets of genetic variation within *E. gracilipes* as separate species. While there is a clear rationale for this proposition, we believe there are reasons for considering an alternative approach. Firstly, there is insufficient evidence that the *E. gracilipes* population groups are independently evolving lineages, given the shallowness of many branches on the phylogenomic trees and degree of nuclear gene tree conflict first indicated by White and colleagues (2021) and supported by gene flow patterns detected in this study. As a second argument, expressed phenotypes remain highly relevant to botanical classification (Wells et al., 2022) not least due to their applicability in ecological contexts such as functional diversity (Sultan, 2000). Rigorous phenotypic analysis, facilitated by advances in the field of morphometrics (Christodoulou et al., 2020) here supports a high degree of consistency in terms of the large, mucronate to acuminate leaves of *E. gracilipes*. Combined with dependable characters used for taxonomy mainly based on lanceolate and coriaceous leaves, and free styles in brevistylar and longistylar flowers, this still supports the retention of a single species. Splitting of *E. gracilipes* could also have consequences for interpretation of the evolution of the domesticated *E. novogranatense* and *E. coca* lineages, reinforcing the idea of domestication events isolated in space and time. Such a model is incongruent with the theory of a protracted timescale of human selection on coca that spans a variety of habitats (Plowman, 1984), now well-supported by the landscape-level domestication paradigm, built on evidence of extensive human-mediated genetic exchange (Allaby et al., 2022; Fuller et al., 2022).

We deduce that the gene pool ancestral to *E. gracilipes* encompassed rich standing genetic variation, from which lineages of *E. novogranatense*, *E. cataractarum*, and *E. coca* var. *coca* also emerged, probably in response to novel environmental and ecological conditions. Our molecular dating results showing clade divergence of the cultivated lineages in the mid to late Pleistocene suggest that that these had already undergone adaptation to ecological conditions distinct from those of their sister wild-living, forest-dwelling relatives; at the point of being selected, these populations were feasibly pre-adapted to a human environment. The species *E. cataractarum* was mostly shaped by adaptation to drier, high-altitude habitats of the eastern Andean slopes (White et al., 2021). Despite limited sampling, genomic data support monophyly of this group. There is contrasting evidence for cocaine in different specimens of *E. cataractarum* (Plowman & Rivier, 1983) and some ethnobotanical evidence of its use (Schultes, 1981), including the inscription on the K type specimen itself ‘Ipadú das Cachocinas’ (‘ipadu’ being an Indigenous name for coca widely used in Brazil; Plowman, 1979). *E. cataractarum* is not documented as a cultivated variety of coca, but since high levels of cocaine are one of the coca’s defining phenotypic traits (Plowman & Rivier 1983), and considering the high degree of leaf morphological overlap with cultivated cocas, it might be considered an intermediate form between wild and domesticated coca.

*E. novogranatense* and *E. coca* var. *coca* have been taxonomically proposed as ‘neo-species’ resulting from long-term human cultivation and selection (Rury, 1981), and the population genomic results confirm the distinctness of these taxa *sensu lato*. We urge caution in labelling these ‘independently evolving monophyletic lineages’ (White et al., 2021), since this not supported by the evidence for gene flow we detected across datasets and overlooks the diffuse nature of domestication (Allaby et al., 2008; Gross & Olsen, 2010). On the other hand, we agree that the differentiation is appreciable, phylogenomically supported, and one argument for retaining their respective species designations. The differentiation could have been driven in part by breeding system: the majority of *Erythroxylum* spp. are reportedly heterostylous (White et al., 2019), with *E. coca* var. *coca* and *E. novogranatense* exhibiting self-incompatibility (Ganders, 1979). Heterostyly has previously been linked to elevated genetic differentiation between neotropical species of *Erythroxylum* (Abarca et al., 2008). In contrast, *E. coca* var. *ipadu*, which is the only variety of coca that is predominantly out- crossing (Plowman, 1979) exhibits genetic identity linked to that of the isotype-defined *E. gracilipes* group, reinforcing our phylogenomic conclusion – of a close genetic relationship of *E. coca* var. *ipadu* to this population of the wild *E. gracilipes.* This puts into question its validity as a variety of *E. coca*.

Our leaf morphometric dataset supplies evolutionary insights which complement the phylogenomics in terms of paraphyletic emergence of coca as a leaf crop. Our most significant finding in this context is that of leaf size distinctness between wild *E. gracilipes* and smaller-leaved domesticated *E. coca* and *E. novogranatense*. *E. coca* var. *ipadu* is not excluded from this pattern, despite its close genetic affinity to *E. gracilipes.* Our data also show a tendency for rounder leaves in *E. coca* compared to wild species. Hence, we provide statistical evidence which to an extent supports cultivated coca leaves being smaller and rounder than those of their wild progenitor (White et al., 2021), a hypothesis built upon in-depth physiological studies of stomatal density and vein morphology (Rury, 1981). The pattern we observe here in the cultivated cocas could be the result of a novel, open habitat of which one facet is greater exposure to sunlight; the domesticates invest fewer resources into leaf expansion than they would in the moist forest habitat of *E. gracilipes*. Experimental work has revealed latent plasticity in leaf size within cultivated coca – manifestation of smaller ‘sun leaves’ and larger ‘shade leaves’ not only in their native Neotropical setting, but also when cultivated under glass at temperate latitudes (Rury & Plowman, 1983). Leaf morphology is generally evolutionarily labile in plants and thus readily adaptable to new microhabitats or biomes (Spriggs et al., 2018). We propose that the consistently smaller leaves in cultivated cocas are the result of genetic canalisation (Flatt, 2005; Piperno, 2017), whereby phenotypic variation in the cultivated species has become to a large extent developmentally constrained to small leaf size. A significant shift in environmental conditions, including escape from human cultivation, may over time unlock persistent cryptic genetic variation (Flatt, 2005), here responsible for the large-leaved *E. gracilipes* phenotype.

*E. cataractarum* is a highly valuable control taxon to benchmark this theory of leaf shape evolution. Having presumably circumvented the selective pressures associated with human cultivation, it has nonetheless attained a leaf size statistically indistinguishable from that of cultivated cocas.

A further possible domestication syndrome trait is acuteness at the leaf base (obovate shape), which contrasts to the frequently observed apical acuteness of wild *E. gracilipes*.

Given the sheer volume of leaves harvested by certain Indigenous communities to supply their daily needs of almost constant coca chewing (Schultes, 1981), it is plausible that reorganisation of the leaf structure to be amenable to human leaf picking could have given these coca bushes an adaptive advantage. Interestingly, the EFA distils basal acuteness as a trait occurring only in the cultivated species and not *E. cataractarum.* Finally, length-to-width ratio of coca leaves could warrant further study; this has for example. been pinpointed as the primary source of variation between accessions and species of apple, with a heritable basis (Migicovsky et al. 2018).

Rury (1981) underscored the challenging, subtle nature of morphological differences between cultivated cocas and their closest wild relatives, which supports the failure of our allometric and EFA analyses to reliably classify coca leaves by current taxonomy. Previous attempts to discriminate Colombian coca cultigens by leaf morphology have been similarly inconclusive (Galindo Bonilla & Fernández-Alonso, 2010; Rodríguez Zapata, 2015). Studies of other plants cultivated for their fruit (Chitwood et al., 2013; Chitwood & Otoni, 2017; Klein et al., 2017) or roots (Gupta et al., 2020) have shown that leaf shape, as inferred through an EFA framework, is highly heritable. EFA has been used to demonstrate that morphological groupings are consistent with species boundaries (Andrade et al., 2008; Klein et al., 2017; Sayıncı et al., 2015), but this is not consistently true across plant groups, particularly those occupying ecosystems more diverse than that of a modern agricultural setting (Nascimento et al., 2021; Soares et al., 2011), in line with the situation for Indigenously cultivated coca.

## Conclusion

Synthesizing the evidence from phylogenomics, gene flow analyses, phenotypic plasticity in coca leaf shape, and the limited discriminatory power of other characters such as leaf venation and floral anatomy (Plowman 1979; Rury 1981), a phylogenetic species concept could be applied here to lump *E. gracilipes*, *E. coca*, *E. novogranatense* and *E. cataractarum* into a single species. Nevertheless, the insights from population genomics and molecular dating do support the identity of long-recognised, and distinct cocas. As such, we propose that the names of these are retained for the time being, so as not to compromise their cultural significance. A future prospect could entail re-classifying *E. c.* var *ipadu, E. c.* var *coca, E. n.* var *novogranatense* and *E. n.* var *truxillense* as varieties of equal standing within a more complex species, to more easily accommodate any new varieties of coca that are described using genomic, ethnobotanical, and metabolomic evidence. We hope that the evidence base we developed will encourage the development of appropriately sensitive genomic methods that can discriminate high-yielding trafficked varieties from varieties used by traditional Indigenous communities in non-harmful ways.

## Materials and Methods

### Leaf morphometrics

#### Image preparation

A total of 1,181 leaf outlines were extracted from 342 digital herbarium specimens, representing individuals collected in the wild (species: *E. gracilipes*, *E. cataractarum*, *E. lineolatum* and *E. foetidum*), as well as plants cultivated in the Neotropics or under glass at temperate latitudes (species: *E. coca* and *E. novogranatense*) (Table S1). Species identities of the specimens were recorded from herbarium labels. We followed this with morphological taxonomic verification of each specimen, based on which we excluded those that could not be confidently assigned to any *Erythroxylum* species of interest to this study (*n*=34) and re- assigned the taxon for several cultivated samples (*n*=14). Ideally, an equal number of leaves would be sampled from each specimen, but sampling was limited by the number of leaves on each specimen that were both intact and fully developed. We set an upper limit of five leaves per specimen. A leaf was considered fully developed if it was roughly the same size as the largest leaf on the sheet and showed shape characteristics consistent with botanical descriptions. Leaves were sampled only if they were pressed flat and intact at the base and apex.

Image features that obstructed leaf contours (e.g., herbarium tape, twigs, and cracks) were manually removed with a digital paintbrush and images were segmented to isolate selected leaves. Outlines were extracted as coordinates using the ‘DiaOutline’software (Wishkerman & Hamilton, 2018). Digital noise was removed from the outlines using the ‘coo.smooth’ function in the R package ‘Momocs’ v1.3 (R Core Team, 2020; Bonhomme et al., 2014). A total of 200 equidistant, non-homologous pseudo-landmarks were sampled from each outline (‘coo.sample’ function in ‘Momocs’; Bonhomme et al., 2014). Two additional landmarks were manually defined at homologous positions on the base and apex of every leaf (‘def_ldk’ function in ‘Momocs’; Bonhomme et al., 2014).

#### Calculating linear metrics and shape variables

Outlines were standardised to 100 pixels per cm and linear metrics (length, width, and area) were recorded for each leaf using the ‘coo_toolbox’ in ‘Momocs’ (Bonhomme et al., 2014). To generate quantitative shape variables, leaf outlines were decomposed by elliptical Fourier analysis (‘efourier’ function in ‘Momocs’; Kuhl & Giardina, 1982; Bonhomme et al., 2014). This type of outline analysis relies on the principle of Fourier series to express an outline as the sum of simpler trigonometric functions. The frequencies of each harmonic in the series are described by four scalar coefficients. These coefficients can be treated statistically as homologous, quantitative variables (Claude, 2008; Zelditch et al., 2004). The minimum number of harmonics necessary for ample shape reconstruction was estimated through estimation of harmonic power (Lestrel, 1997; Bonhomme et al., 2014). We computed 14 harmonics, representing a cumulative harmonic power near 100%. Because the coefficients of an elliptic Fourier descriptor are not invariant in size, rotation, shift and starting point of chain-coding about a contour (Yoshioka et al., 2004), we standardized the Fourier coefficients prior to this shape analysis. Outlines were normalised with Bookstein baseline superimposition to the two homologous landmarks defined in outline preparation (‘fgProcrustes’ function in ‘Momocs’; Friess & Baylac, 2003; Bonhomme et al., 2014). The dataset was passed through a second round of normalisation, as part of the default ‘efourier’ function in ‘Momocs’.

Leaves with aberrant shapes heavily skew dimensionality reduction, thus we removed these from the dataset prior to further analyses. Aberrant leaf shape outliers were identified by modelling the shape variation for each species as normally distributed, with a confidence level of 1e^-3^ (‘which_out’ function in ‘Momocs’; Bonhomme et al., 2014). In total, 15 leaves with aberrant shapes were excluded. Upon inspection, aberrant contours were mostly due to distortion from the drying process.

#### Morphometrical statistical analyses

Shape and size are separate aspects of an organism’s morphology, and natural patterns can be obscured if an investigation should confound the two (Christodoulou et al., 2020). Therefore, we treated linear metrics and shape variables separately in statistical analyses. To define morphological spaces for respective linear metric and shape cluster analyses, data dimensionality was reduced via principal component analysis (PCA) (Claude, 2008). To visualise the distribution of the most common leaf shapes within a species, we performed PCA separately for each taxonomic subset of Fourier data, after which eigen-leaves were reconstructed using the reverse Fourier transform for the mean, and +/- 1.5 standard deviations from the mean, on the first two PCs of shape variation within a species (‘PCcontrib’ function in ‘Momocs’; Bonhomme et al., 2014). To assess the distribution of our taxon-binned leaf data along the first two axes of variation, corresponding to defining leaf traits, we constructed eigen- leaves for the first two PCs of the shape (EFA) PCA, encompassing 95.4% of the variation combined.

To investigate the question of whether underlying structure in this morphometric dataset reflects current taxonomic boundaries, variation within this dataset was examined without *a priori* identifications. We inferred morphological groups probabilistically using model-based clustering using Gaussian mixture models (GMMs; Bouveyron et al., 2019; Bouveyron & Brunet-Saumard, 2014). This method of cluster analysis assumes that continuously distributed data can be described as a mixture (weighted average) of *G* multivariate normal distributions (clusters). The aim is to identify the number of groups (*G*) and the geometric properties of their densities; different model parameters correspond to different hypotheses of group structure (Bouveyron et al., 2019). Model selection was performed with the expectation-maximization (EM) algorithm, which estimates model parameters by maximum likelihood (Dempster, et al., 1977). Measures of empirical support are based on the Bayesian Information Criterion (BIC; (Schwarz, 2007), wherein Δ BIC expresses the gain in the explanatory power of the model when an additional group is considered by the algorithm. Based on Δ BIC, an EEV geometrically-constrained clustering model (ellipsoidal, equal volume and shape) was selected for Fourier data and an EVE model (ellipsoidal, equal volume and orientation) was selected for linear metrics. Each leaf was assigned to a cluster by soft classifications – expectations of the assignments under the applied probability model. GMMs were fitted using the ‘mclust’ v5.0 package (Scrucca et al., 2016), with parameters set to model two to fifteen groups. If leaves are well-classified by a given model, the conditional probability that a leaf belongs to one of the postulated groups should be close to 1. To statistically quantify equivalence between taxonomic clusters and GMM-inferred clusters, we applied the Rand index, whose value indicates the degree of correspondence of two different clustering outcomes from 0 = complete mismatch 1 = to perfect match (Rand, 1971). We also applied the adjusted Rand index which corrects for chance (Hubert and Arabie, 1985), carrying out all analyses in the ‘Fossil’ package (Vavrek, 2011).

Noisy variables in high-dimensional data are known to degrade the performance of model-based clustering, so variable selection is crucial (Bouveyron et al., 2019). Only three morphological variables were recorded for leaf size, so data were relatively low-dimensional. As such, all three principal components (PC) of their log-transformation were used in cluster analysis. Although Fourier descriptors of leaf shape are more complex in dimension, experimental exploration of a modelling approach to model selection (Maugis et al., 2009) did not provide valuable insight. Therefore, we retained the 4 PCs explaining most of the variation, deeming these most useful in defining group structure following previous studies (Ezard et al., 2010; Sneath & Sokal, 1975). To visualise clustering for each leaf on a given herbarium specimen, bar charts were created to represent the number of leaves sampled, and their individual cluster assignments (Fig S1).

#### Canonical variants analysis

To visualise relationships between the *a priori* taxonomically defined groups in morphological space, we also computed mean vectors for the shape variables and Euclidean distances between these were calculated using the R package ‘rdist’ (Blaser, 2020) and the output distance matrix was used for Principal Coordinates Analysis (PcoA) (‘pcoa’ implemented in R package ‘ape’).

#### Dataset filtering and downsampling

Finally, we carried out four iterations of filtering our full datasets of digitalised leaf outlines. The primary purpose was to produce a dataset excluding those samples assigned dubious IDs after the taxonomic evaluation and those which were cultivated outside of South America. This dataset consisted of 856 leaves from 260 specimens. We reran the analyses using the filtered dataset, using this dataset for our main interpretation of the results. The second filtering iteration had the goal of demonstrating that there was no bias due to different sample sizes for different taxa. Therefore, for each pre-assigned taxonomic group, we downsampled to a random subset 50 leaves (excluding *E. lineolatum*, for which this quota was not reached). The final two iterations involved retaining from our main interpretation dataset only the cultivated coca species with a main goal of computing the degree of correspondence between the taxonomic groups and GMM clusters. One of the iterations comprised the full dataset of *E. coca* and *E. novogranatense* individuals (404 samples total) and the other is a subset of these, including only one leaf per specimen (97 samples total).

### Genomics

#### Taxon sampling and genomic data mining

Our taxon sampling builds upon previous phylogenomic and population genomic studies of the coca clade (White et al., 2019; 2021). We generated data from 20 accessions of coca and its closest wild relatives following the current taxonomy (Plowman, 1982; White et al., 2019, 2021). Our sampling included 12 specimens of *E. gracilipes* (including a Kew [K] isotype), 3 of *E. cataractarum* (including a K holotype), 1 of *E. lineolatum* and 1 of *E. foetidum* from K as well as 1 specimen of *E. n. novogranatense* (S.S. Renner 2888) (Table S2). For each sample, we weighed out 0.2-0.3g of leaf tissue and pulverised this using steel ball bearings in a SPEX® sample prep tissue homogeniser (SPEX Inc, NJ, USA). We extracted genomic DNA following a CTAB protocol, with the addition of 2% v/v 2- mercaptoethanol (Doyle & Doyle, 1990). DNA extracts were purified using a bead clean up method with a 2:1 ratio of Ampure XP beads (Beckman Coulter, USA) to DNA elution. DNA extracts were quantified using a Quantus® fluorometer (Promega, USA) and the degree of DNA fragmentation was assessed using a 4200 TapeStation system (Agilent Technologies, USA). We conducted library preparation using a NEBNext Ultra II DNA Library Preparation Kit according to the manufacturer’s protocol. Sequencing of DNA libraries was carried out on an Illumina HiSeq platform with a paired end 150bp configuration, by GeneWiz® (South Plainfield, NJ). To complement this *Erythroxylum* genomic dataset, we mined read data for wild relatives and cultivated species of coca from NCBIs Sequence Read Archive (SRA) repository. This comprising raw sequence data from 155 herbarium specimens from which DNA had been extracted and enriched for 427 nuclear genes via target capture with a custom-designed set of RNA probes (White et al., 2019; 2021).

#### Processing of raw read data and alignment to target sequence

We trimmed raw reads (both sequenced and data-mined) using AdapterRemoval v2.1 (Schubert et al., 2016). With the goal of retrieving nuclear genomic information, the trimmed reads were mapped against a set of low-copy nuclear genes selected to infer phylogenetic relationships in *Erythroxylum* (White et al., 2019), totalling 419 genes after having removed those flagged as potential paralogs. The reads were mapped using Bowtie v.2.3.4.1., after which they were realigned around indels using GATK v.3.8.1 and filtered for duplicates using picard-tools, all steps being executed within the pipeline ‘PALEOMIX’ v.1.2.13 (Schubert et al., 2014). Similarly, the data were mapped to a published *E. novogranatense* plastid genome from NCBI using the PALEOMIX pipeline and settings identical to those above.

#### Plastid and nuclear phylogenomic analysis

To investigate phylogenetic relationships between recognised wild relatives and cultivated taxa of coca, we generated sequence data from reads mapped both to the full plastid genome and to the low-copy nuclear genes. Our primary aim was to produce, for the first time, a plastid phylogeny for this clade. We opted for a pseudohaploid approach to sequence generation owing to the low read depth across the nuclear genes – a consequence of our genome skimming approach. Using our data from 17 samples mapped to the *E. novogranatense* plastome, we generated pseudohaploid sequences using ANGSD v. 0.930 (Korneliussen et al., 2014) sampling a base at random (with option -doFasta 1), setting a minimum mapping quality of 25, a minimum base quality of 25 and minimum depth of 10.

From the published datasets, we successfully data-mined plastomes of 9 samples, and generated sequences from these in the same manner, but requiring a min. depth of 3 to retain a site and genome completeness of min. 90% retain a sample. This low rate of ‘by-catch’ plastome recovery from the data-mined reads was due to the authors’ experimental aim of nuclear gene target enrichment. In addition, we downloaded an *E. novogranatense* plastome from NCBI to use as a reference. We aligned to this reference our dataset of 25 samples using MAFFT v7.508 with the fast, progressive strategy (FFT-NS-2), which applies two guide tree builds and a maximum of 1000 iterations. To build a phylogenetic representation of relationships between the samples, we ran RAxML v. 8.2.12 (Stamatakis, 2014) under the rapid bootstrap analysis mode, with the GTR substitution model, the GAMMA model of rate heterogeneity and 500 bootstrap iterations.

To build a nuclear phylogeny that would correspond to our plastid phylogeny, we aimed for identical sampling. We used our dataset of K samples aligned to the low copy nuclear genes (see *Processing of raw read data and alignment to target sequence*), where 14 of the 17 samples yielded sufficient data for inclusion. We also included data-mined samples of *E. foetidum*, *E. gracilipes*, *E. cataractarum*, *E. novogranatense* and *E. coca* from which we had retrieved ‘by-catch’ plastid data (Table S3) and used in the plastid tree. Finally, we data-mined accessions of *E. lineolatum* (*n*=*1*) and *E. williamsii* (*n*=*1*) from the NCBI repository as outgroups for the study (based on White et al., 2019). Notably, a different sample of *E. foetidum* was used in the nuclear phylogeny, due to insufficient retrieval of nuclear genes from our genome skimmed *E. foetidum* 13512 sample. These different samples used are in fact geographically proximate; *E. foetidum* 13512 is from the Átures Department in Venezuela and *E. foetidum* F_2324334 is from the Vichada Department of Colombia.

For all alignments, we generated pseudohaploid sequences, by sampling a random allele at each site using ANGSD, requiring a minimum depth of 3, a minimum quality score of 25 and minimum mapping quality of 25. We took forward for phylogenomic analysis those samples for which at least 134 genes were genotyped to at least 80% of their total length.

Two samples were retained as outgroups - one *E. foetidum* sample as a first outgroup, and one *E. williamsii* as the second outgroup, seeing that no *E. lineolatum* samples passed the ‘minimum number of genes’ threshold. A total of 326 genes, qualifying based on their retention in at least 75% of the samples, were used in the final alignment. For each gene, we computed a gene tree using RAxML v.8.2.12 with the rapid bootstrap analysis mode, a GTR+GAMMA model of substitution and 500 bootstrap replicates. The resulting 326 gene trees were summarised into a multispecies coalescent tree phylogeny using gene-tree reconciliation in ASTRAL-III v.5.7.8 (Zhang et al., 2018). We did not collapse any of the branches since the lowest posterior probability support value was 0.4. To assess incongruence between the 326 gene trees used to make the species tree, we began by conducting a bipartition analysis with PhyParts (Smith et al., 2018). This method, which leverages added precision gained by using rooted gene trees as input, was used to summarise at each node the conflicting, concordant and unique bipartitions with respect to the ASTRAL species tree topology. We visualised the output using a script that plots pie charts (Johnson, 2021).

To further interrogate gene tree support for species tree, we computed metrics which have complementary explanatory power: gene concordance factors (gCF; Baum, 2007) and internode certainty (IC, ICA). For every bipartition of the tree, the gene concordance factor is the percentage of decisive trees that contain that bipartition (Minh et al., 2020), whereas IC is a quantified degree of certainty for individual bipartitions which considers the frequencies of the most frequent specifically conflicting bipartition (Salichos & Rokas, 2013). ICA in turn considers all gene trees with a decreasing logarithmic weight (Salichos et al., 2014). We computed these metrics using a custom script designed to consider well-supported bipartitions on the basis of gene tree bootstrap values in gene trees with variable support among branches (Simon et al., 2022). Finally, using fasta alignments, we also computed site concordance factors (sCF; Minh et al., 2020), which measure the percentage of decisive sites supporting a branch in the reference tree. This was done using IQ-TREE2 (Minh et al., 2020).

As a supplementary exploration of the nuclear phylogeny, we concatenated alignments of the 326 genes and constructed a maximum likelihood (ML) tree using RAxML v. 8.2.12 with the GTRCAT model of substitution, to produce a reference tree against which we interrogated the ML gene trees. We also computed support metrics gCF, IC and ICA for this tree using the same method as above. Finally, we plotted all output trees in FigTree v.1.4.4 (Rambaut, 2016).

#### Molecular clock dating analyses

The time of origin and diversification of the tropical American lineages of *Erythroxylum* remain elusive. The first and only inference of absolute ages of *Erythroxylum* was conducted by White (2019), who relied on a penalised likelihood approach implemented on a phylogram derived from a concatenated supermatrix of 544 nuclear genes plus a fossil constraint applied to the stem node of the stem lineage of the tropical American *Erythroxylum* and secondary calibrations. However, the cultivated cocas were not included in this analysis, and *E. williamsii*, a taxon deemed sister to *E. lineolatum* and the coca clade by White and colleagues (2019), shared a common ancestor with *E. cataractarum* and *E. gracilipes* ∼20 Ma.

Because estimation of absolute ages based on concatenated supermatrices are known to be prone to produce erroneous branch lengths in the presence of incomplete lineage sorting (Ogilvie et al., 2017), here we opted to derive ages of divergence using a Bayesian Multi-species Coalescent approach that is known to reliably estimate calibrated time trees, even in the presence of gene tree conflict. This approach uses as input multiple gene alignments, age prior calibrations, and prior knowledge of population memberships (Ogilvie et al., 2017; Yan et al., 2022). To obtain ages of divergence in the coca clade, we first estimated the root-to-tip tree variance, concordance and length of our 326 Maximum Likelihood gene trees (the same as input into species tree inference through gene tree reconciliation in ASTRAL – see *Plastid and nuclear phylogenomic analysis*), using SortaDate v.1.0 (Smith et al., 2018). We then selected the 20 most clock-like (i.e., lowest root-to-tip variance), congruent gene alignments that represented the entire panel of samples and populations included in the ASTRAL-III species tree analysis. The dataset subset as such was imported into BEAUTi v.2.6 (Bouckaert et al., 2019) as unlinked partitions using the following priors *a*) a weakly informative secondary calibration applied to the root of the tree (i.e., the MRCA of *E. williamsii*, *E. lineolatum* and the coca clade) modelled by an exponential distribution with lambda equal to 0.1498, so that 95% of the density is between 0 and 20 Ma, and the mean 6.60 Ma; b) an uncorrelated log-normal relaxed clock with a lognormal prior on the mean rate with 95% of the density between 0.0001-0.001 substitutions/site/Ma with parameters log-mean -6.909 and log-standard-deviation 1.1746 c) a ploidy level of 2 (option “autosomal nuclear”) as recommended for diploid organisms and following the known ploidy levels of the cultivated cocas (Rodríguez-Zapata, 2015); d) a Coalescent Constant Population tree model with a mean population size of 1.0 and a non-informative prior of 1/X (as recommended by Drummond & Bouckaert [2015] for datasets that contain population level samplings); e) a theoretical number of populations (*K*) of four, following the clusters produced by NGSadmix and that attained a high delta Likelihood. We used the function findParams of the tbea package (Ballen & Reinales, 2021) to find the parameter values that best describe the probabilistic expectations in the priors. We ran the molecular clock analyses for 1 billion generations, sampling every 50,000 states and ensuring that all parameters converged by attained effective sample sizes > 200. Additionally, we ran three independent analyses and examined the posterior marginal distribution for each parameter, to check whether convergence was attained beyond within-change convergence measures such as ESS. We found that the three independent runs arrived at essentially the same posterior distributions for all the parameters in the model. Lastly, to ensure that all priors implemented were informative, we conducted one independent analysis where sampling was drawn from the priors, which revealed that indeed, our prior parameters were informative. Parameter and tree summaries were generated and visualised using Tracer v1.7.2 (Rambaut & Drummond, 2013) and Treeannotator v.1.0 (available at https://beast.community/tree-annotator).

#### Phylogenetic networks

To explicitly estimate sources and directionality of gene flow within the coca clade, we made use of the 326 loci which comprise the same genes as used in our phylogenomic tree reconstructions (see *Plastid and nuclear phylogenomic analysis*), with which we inferred phylogenetic networks using a pseudolikelihood approach (Solís-Lemus & Ané, 2016). This was implemented in the Julia package PhyloNetworks (Solís-Lemus et al., 2017). The input for SNaQ (Species Networks applying Quartets) normally comprises gene-tree point estimates, however, we chose to use posterior gene tree densities estimated via Bayesian inference to account for topological uncertainty. Gene tree posterior densities were estimated for each locus using MrBayes (Huelsenbeck & Ronquist, 2001). After assessing topological convergence using the standard deviation of split frequencies < 0.05 (SDSF, Nylander et al., 2008), we formatted and thinned the posterior gene-tree sample using phyx (Brown et al., 2017). PhyloNetworks was then used for calculation of concordance factors (Solís-Lemus et al., 2017) using the composite tree sample. Network estimation was carried out using SNaQ (Solıs-Lemus and Ané, 2016) with the same initial tree used in divergence time estimation and considering the number of hybridisations to be h = 0 to 3. Each analysis included 20 independent runs to aid optimisation via more thorough exploration of parameter space. We applied the heuristic criterion of gradient stabilisation to decide how many hybridisations to allow in the network. Finally, a bootstrap analysis was carried out with 100 replicates, each running 20 parallel searches. Credible intervals for the inheritance proportion in minor and major hybrid edges were constructed from the quantiles 0.025 and 0.975 of their bootstrap samples. Phylogenetic networks were plotted using the package PhyloPlots, available at https://github.com/cecileane/PhyloPlots.jl.

#### Population genomic analysis

To examine the shared variance between samples in our dataset of coca wild relatives and to explore patterns of clustering both within this dataset and in the context of published data from wild relatives and cultivated coca, we conducted population genomic analyses based on genotype likelihoods (GLs). We called the GLs using ANGSD v. 0.930 (Korneliussen et al., 2014), setting a minimum p-value of 1e^-6^ to call a variant, a minimum quality score of 25, a minimum mapping quality of 25, and -remove_bads 1 to exclude any possible leftover duplicate or failed reads. We used the matrix of per-sample genotype likelihoods to carry out principal component analysis using PCangsd v.1.02 (Meisner & Albrechtsen, 2018) with 10,000 iterations and requiring a minimum of 50% samples to be genotyped at any site. We also set a minimum minor allele frequency (MAF) of 0.2 (our dataset, *n*=19) and a MAF of 0.05 (merged dataset, *n*=174).

Furthermore, we modelled population structure for the merged dataset using ‘NGSadmix’ v32 (Skotte et al., 2013). We prepared the dataset by filtering for missingness of individuals at a site (tolerating up to 50% missingness) and setting a minor allele frequency (MAF) threshold of 0.05. A total of 13,643 filtered sites were analyzed with default parameters for a maximum of 2,000 expectation maximisation (EM) iterations. We modelled the per-individual ancestries assuming from two to ten ancestral populations (*K* = 2-10), running 10 iterations of the analysis per value of K with different random seeds. We computed the theoretical best value of K, using the Evanno method (Evanno et al., 2005), as implemented in CLUMPAK (Kopelman et al., 2015), identifying the highest values of Δ*K*.

To complement our phylogenomic analysis of plastid data, we also computed a PCA based on variation within the dataset of 17 generated and 8 data-mined coca clade plastids. We ran PCangsd on this dataset, setting a MAF of 0.1, analysing a total of 4,664 sites.

## Data availability

All high-throughput sequencing files are archived in the NCBI Sequence Read Archive (SRA) database under the accession number (*tbc*). All morphometric datasets are available on (*open access data sharing site tbc*).

## Author contributions

OAPE, RAD and AA conceived the study. FAA conducted taxonomic evaluations prior to specimen analysis. RAD conducted morphometric analyses, with mentorship from DC and contributions from NASP. The initial morphometric analyses were conducted by RAD for his MSc thesis (Queen Mary University of London & RBG Kew). NASP conducted wet lab work. NASP, OAPE and LK conducted phylogenomic analyses. NASP and OAPE conducted population genomic analyses. GAB and OAPE carried out divergence time estimation analysis. GAB conducted phylogenetic network analysis. SSR provided samples and advised on taxonomy. RC-B. shared taxonomic expertise and ideas for discussion. OAPE and NASP produced figures, with contributions from RAD and MC.

NASP wrote the manuscript, with contributions from RAD, OAPE and AA. All co-authors read and approved the final manuscript.

## Acknowledgments

We thank Sue Zmartzy from RBG Kew for curatorial assistance with the herbarium specimens and Eve Lucas for consultation on general taxonomic practice and type specimens. We thank the laboratory staff at the Jodrell, and particularly Robyn Cowan for support enabling wet lab work. We are grateful to Carly Cowell for consultation on policy. A subset of computational analyses performed for this paper was conducted on the Smithsonian High Performance Cluster, Smithsonian Institution: https://doi.org/10.25572/SIHPC. AA acknowledges support from the Swedish Research Council (2019-05191), the Swedish Foundation for Strategic Environmental Research MISTRA (Project BioPath), and the Kew Foundation. OAPE is supported by the Sainsbury Orchid Fellowship at the Royal Botanic Gardens Kew and the Swiss Orchid Foundation.

