## Supplementary Materials for "Morphometrics and phylogenomics of coca (*Erythroxylum* spp.) illuminate its reticulate evolution, with implications for taxonomy"

[*]Lead authors

[§]Senior authors

[†]Corresponding authors

1 School of Biological Sciences, University of Portsmouth, Portsmouth, PO1 2DY, UK
2 Royal Botanic Gardens, Kew, Richmond, Surrey, TW9 3AE, UK
3 Department of Anthropology, National Museum of Natural History, Smithsonian Institution, Washington DC, 20560, USA
4 The New York Botanical Garden, New York, NY 10458, USA
5 School of Biological and Behavioural Sciences, Queen Mary University of London, E1 4NS, UK
6 Universidad Distrital Francisco José de Caldas, CR 5E 15-82, Bogotá, Colombia
7 Department of Horticulture and Department of Computational Mathematics, Science & Engineering, Michigan State University, East Lansing, MI 48824, USA
8 Department of Biological Applications and Technology, University of Ioannina, 45110 Ioannina, Greece
9 Department of Biology, Washington University, Saint Louis, MO, 63130, USA
10 Gothenburg Global Biodiversity Centre, Department of Biological and Environmental Sciences, University of Gothenburg, Carl Skottsbergs Gata 22b, se 41319, Göteborg, Sweden
11 Department of Biology, University of Oxford, OX1 3RB, UK

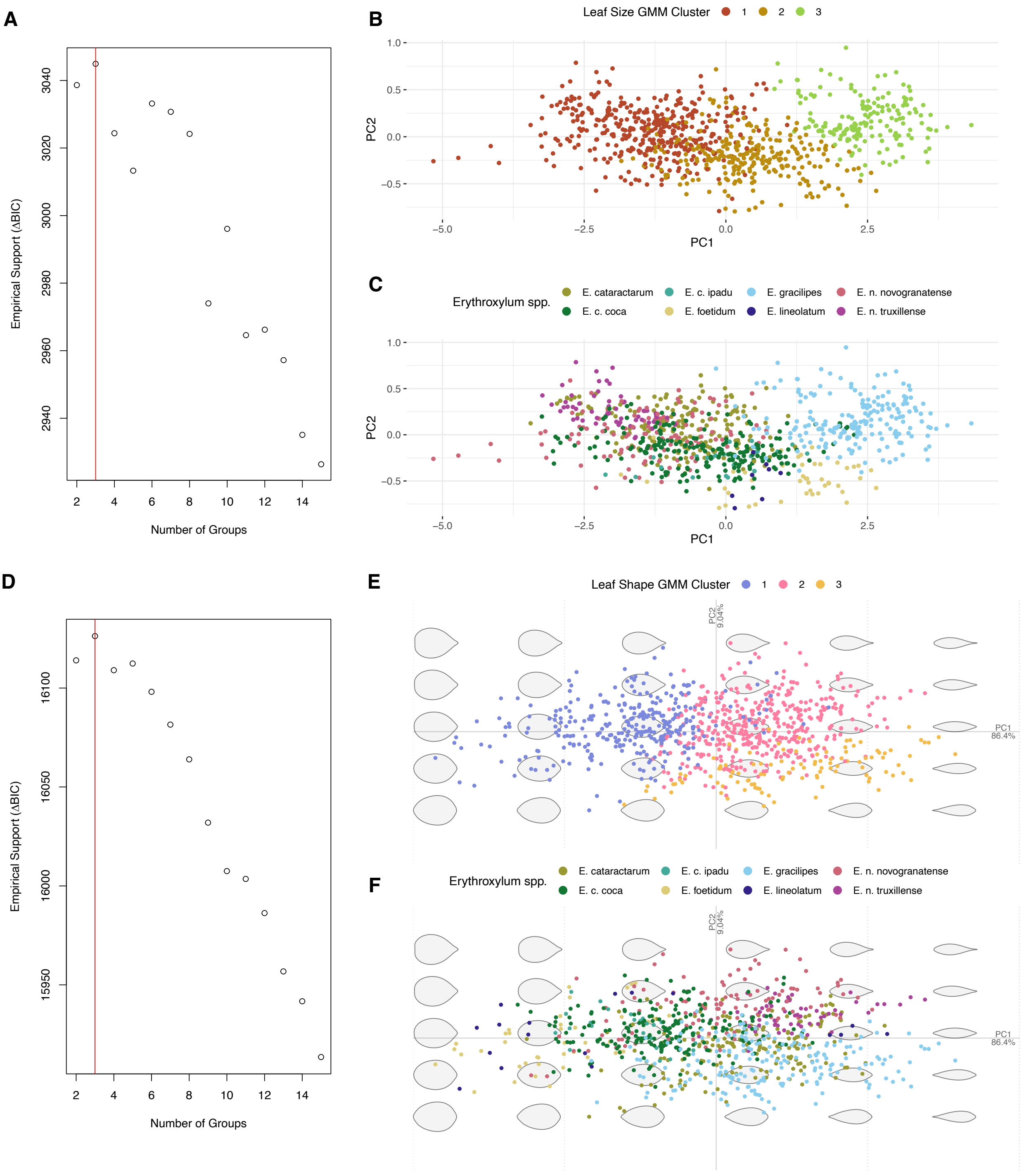


**Figure S1**. **Visual summary of principal component analysis (PCA) conducted with allometric and elliptical Fourier analysis (EFA), grouped by taxonomy or Gaussian Mixture Model (GMM) clusters, and inference of support for number of clusters. Principal dataset (835 leaves total) – constituting all samples except for those with dubious IDs and those cultivated outside of South America or in a glasshouse. (A)** BIC-inferred model with highest support for linear metric data. Highest support for 3 groups indicated. EEI model (diagonal, with equal volume and shape) and variable selection applied. **(B)** PCA of morphospace based on linear metrics. Data coloured by clustering assignments resulting from the GMM. **(C)** PCA of morphospace based on linear metrics. Data coloured by pre-defined taxonomic identifications. **(D)** BIC-inferred model with highest support for EFA data. EEV geometrically-constrained clustering model (ellipsoidal, equal volume and shape) applied. **(E)** PCA of morphospace based on EFA data. Data coloured by clustering assignments resulting from the GMM. **(F)** PCA of morphospace based on EFA data. Data coloured by pre-defined taxonomic identifications.

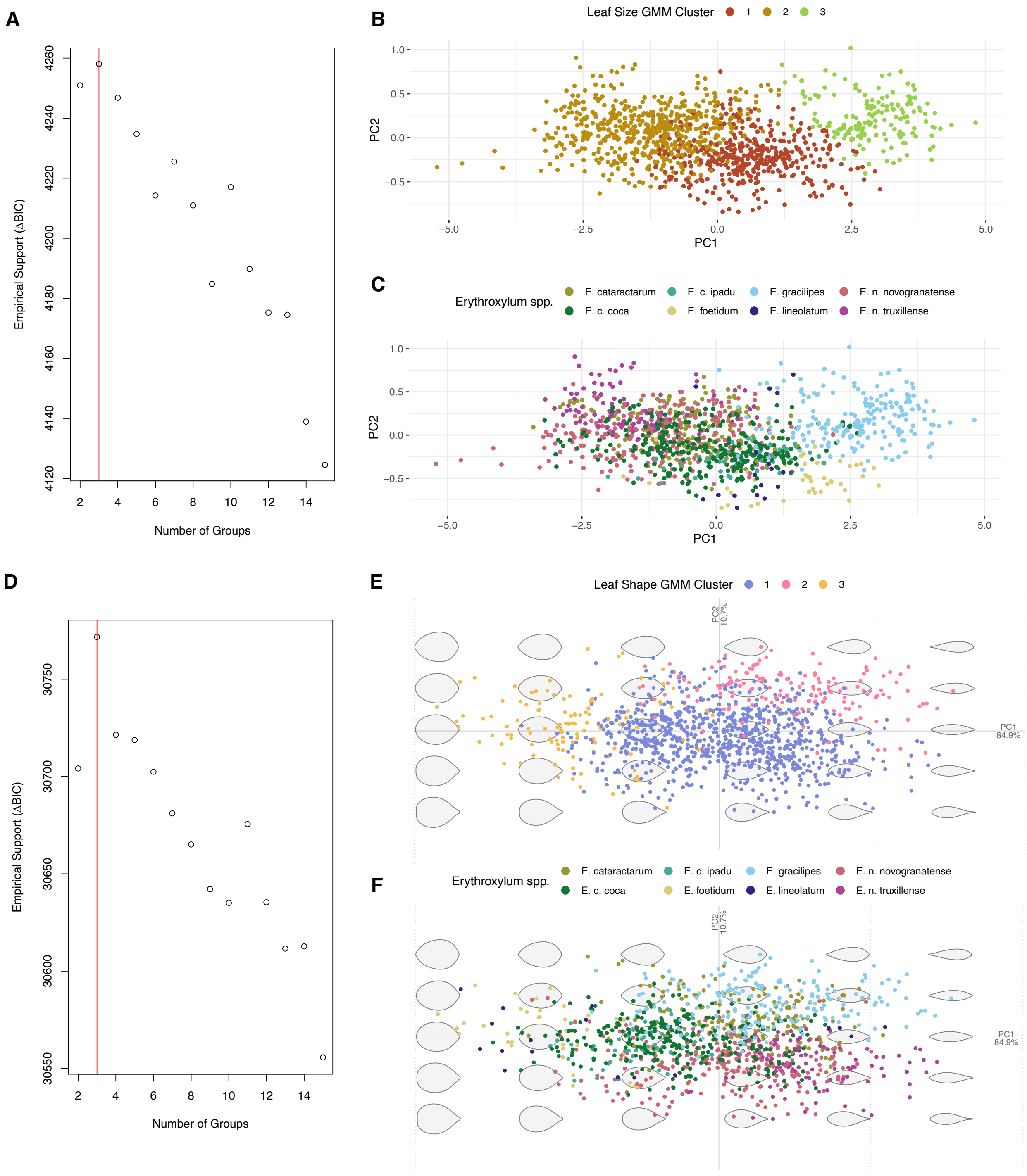


**Figure S2**. **Visual summary of principal component analysis (PCA) conducted with allometric and elliptical Fourier analysis (EFA), grouped by taxonomy and Gaussian Mixture Model (GMM) clusters, and inference of support for number of clusters. Full dataset (1163 leaves total) – including samples with dubious IDs and those cultivated outside of South America or in a glasshouse. (A)** BIC-inferred model with highest support for linear metric data. Highest support for 3 groups indicated. EEI model (diagonal, with equal volume and shape) and variable selection applied. **(B)** PCA of morphospace based on linear metrics. Data coloured by clustering assignments resulting from the GMM. **(C)** PCA of morphospace based on linear metrics. Data coloured by pre-defined taxonomic identifications. **(D)** BIC-inferred model with highest support for EFA data. EEV geometrically-constrained clustering model (ellipsoidal, equal volume and shape) applied. **(E)** PCA of morphospace based on EFA data. Data coloured by clustering assignments resulting from the GMM. **(F)** PCA of morphospace based on EFA data. Data coloured by pre-defined taxonomic identifications.

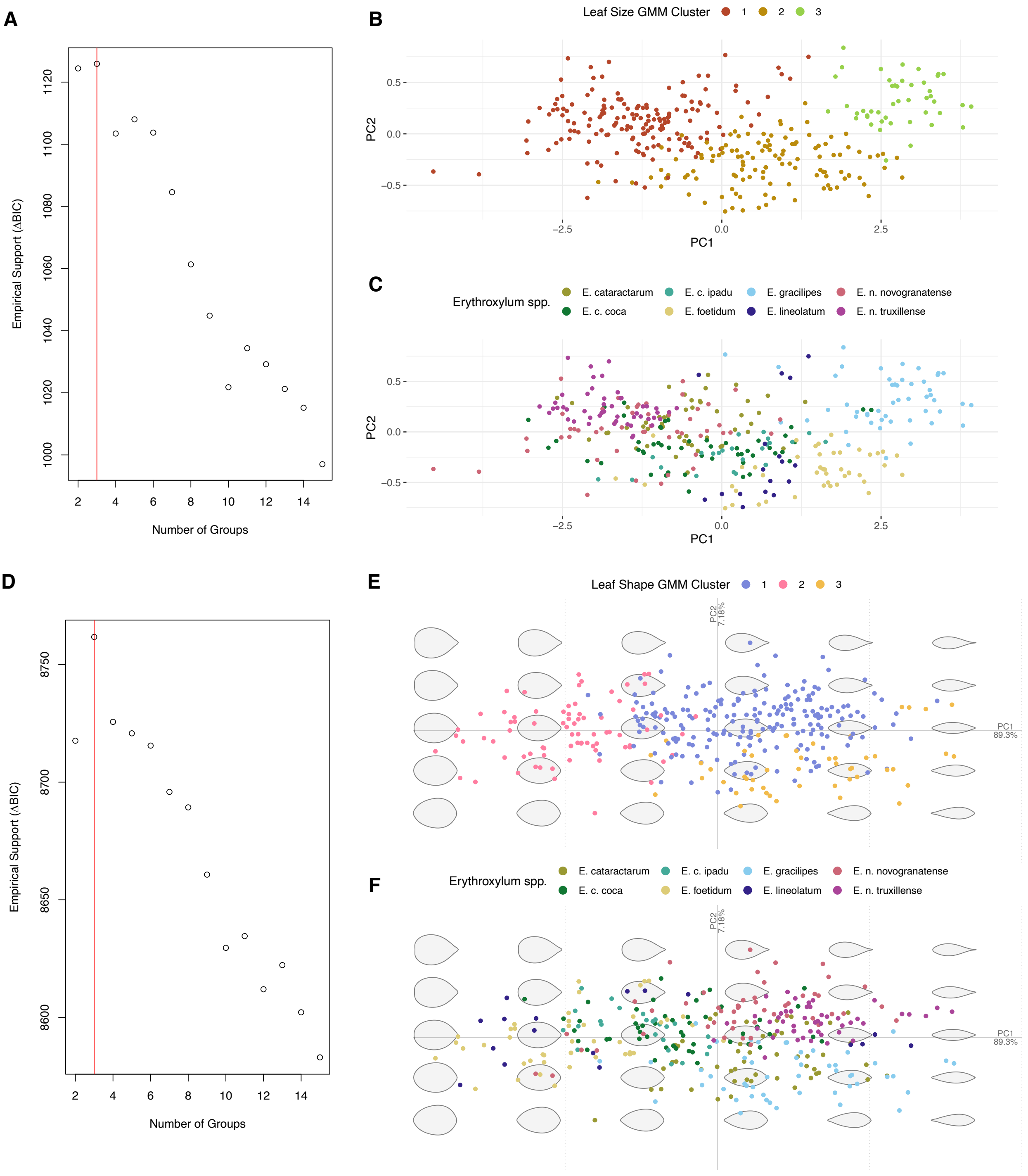


**Figure S3**. **Visual summary of principal component analysis (PCA) conducted with allometric and elliptical Fourier analysis (EFA), grouped by taxonomy and Gaussian Mixture Model (GMM) clusters, and inference of support for number of clusters. Down-sampled database (335 leaves total) – excluding samples with dubious IDs and those cultivated outside of South America or in a glasshouse and with a maximum of 50 leaves per taxon retained.** **(A)** BIC-inferred model with highest support for linear metric data. Highest support for 3 groups indicated. EEI model (diagonal, with equal volume and shape) and variable selection applied. **(B)** PCA of morphospace based on linear metrics. Data coloured by clustering assignments resulting from the GMM. **(C)** Principal component analysis (PCA) of morphospace based on linear metrics. Data coloured by pre-defined taxonomic identifications. **(D)** BIC-inferred model with highest support for EFA data. EEV geometrically-constrained clustering model (ellipsoidal, equal volume and shape) applied. **(E)** PCA of morphospace based on EFA data. Data coloured by clustering assignments resulting from the GMM. **(F)** Principal component analysis (PCA) of morphospace based on EFA data. Data coloured by pre-defined taxonomic identifications.

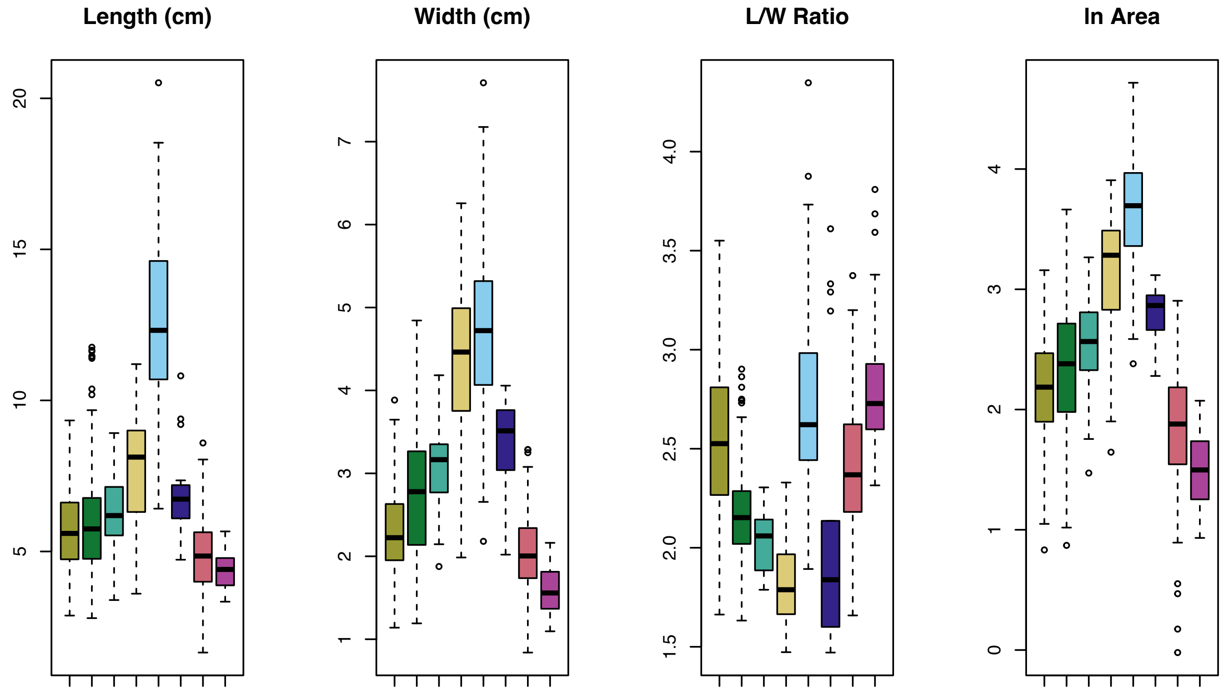


**Figure S4**. Boxplots summarising linear metrics across groups comprising recognised taxa. Summary boxplot from left: length, width, length/width ratio, natural log of area. Within each boxplot from left: *E. cataractarum* (khaki), *E. coca* var. *coca* (dark green), *E. coca* var. *ipadu* (mint green), *E. foetidum* (yellow), *E. gracilipes* (light blue), *E. lineolatum* (dark blue), *E. novogranatense* var. *novogranatense* (pink), *E. novogranatense* var. *truxillense* (fuschia).

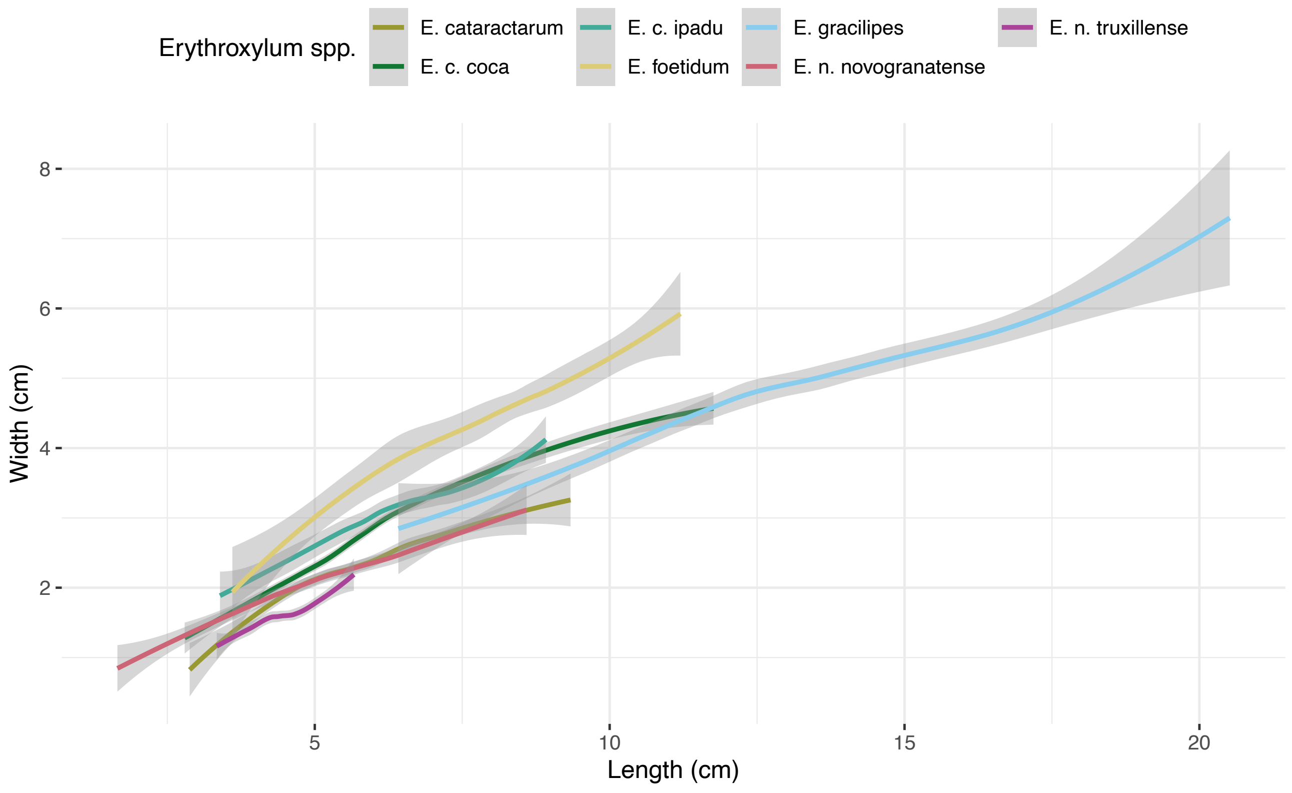


**Figure S5**. LOESS regression curves (95% confidence limit) for allometric relationships of width vs. length, data grouped by pre-defined taxonomic identification.

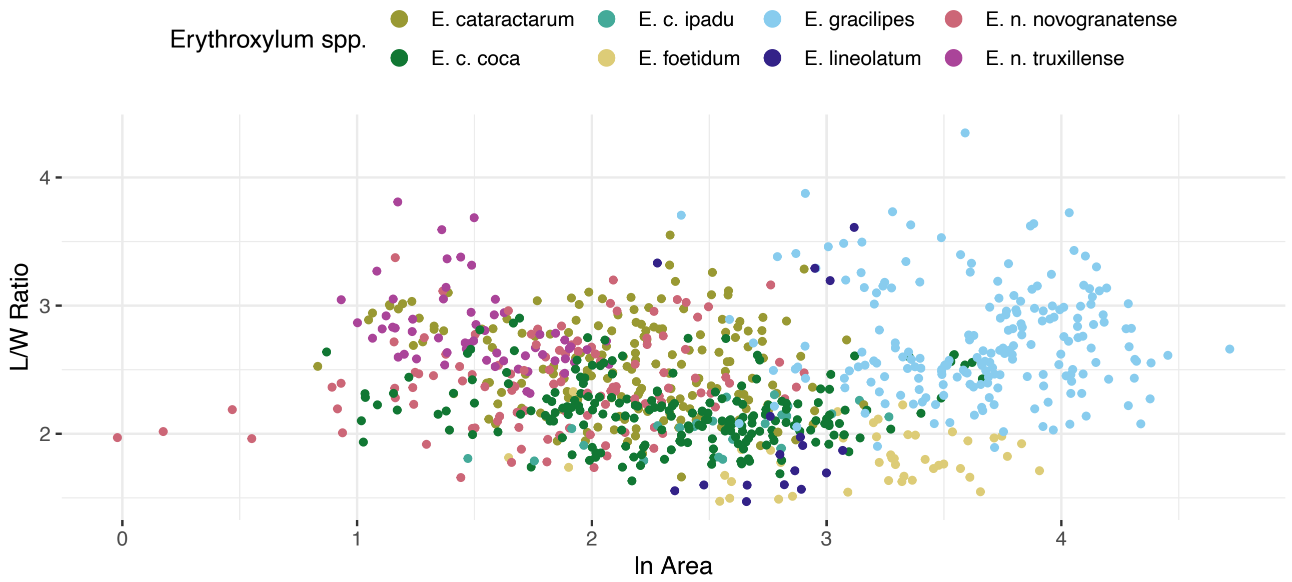


**Figure S6.** Plot of area against length/width ratio. Data coloured by pre-defined taxonomic identification.

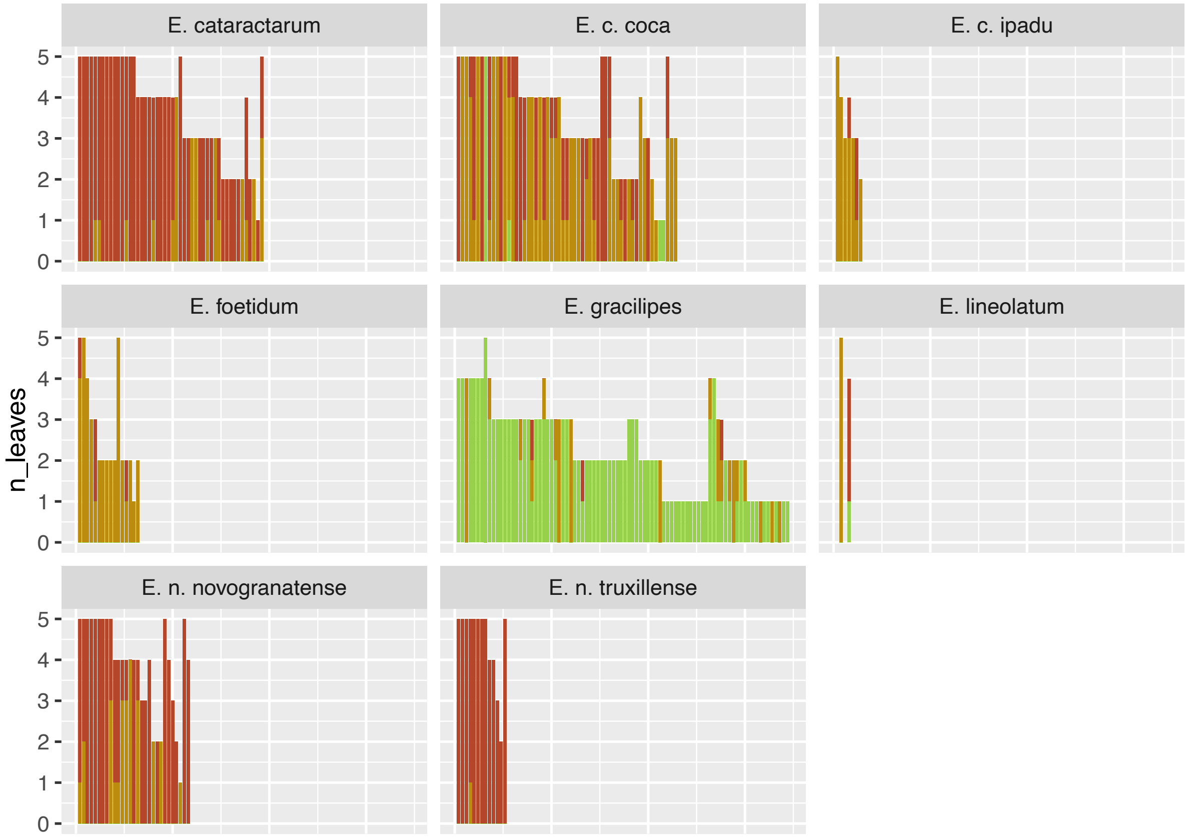


**Figure S7.** Bar charts representing the number of leaves sampled, and their individual cluster assignments based on GMM of allometric data (principal dataset of 835 leaves), and grouped by pre-defined taxonomic identification.

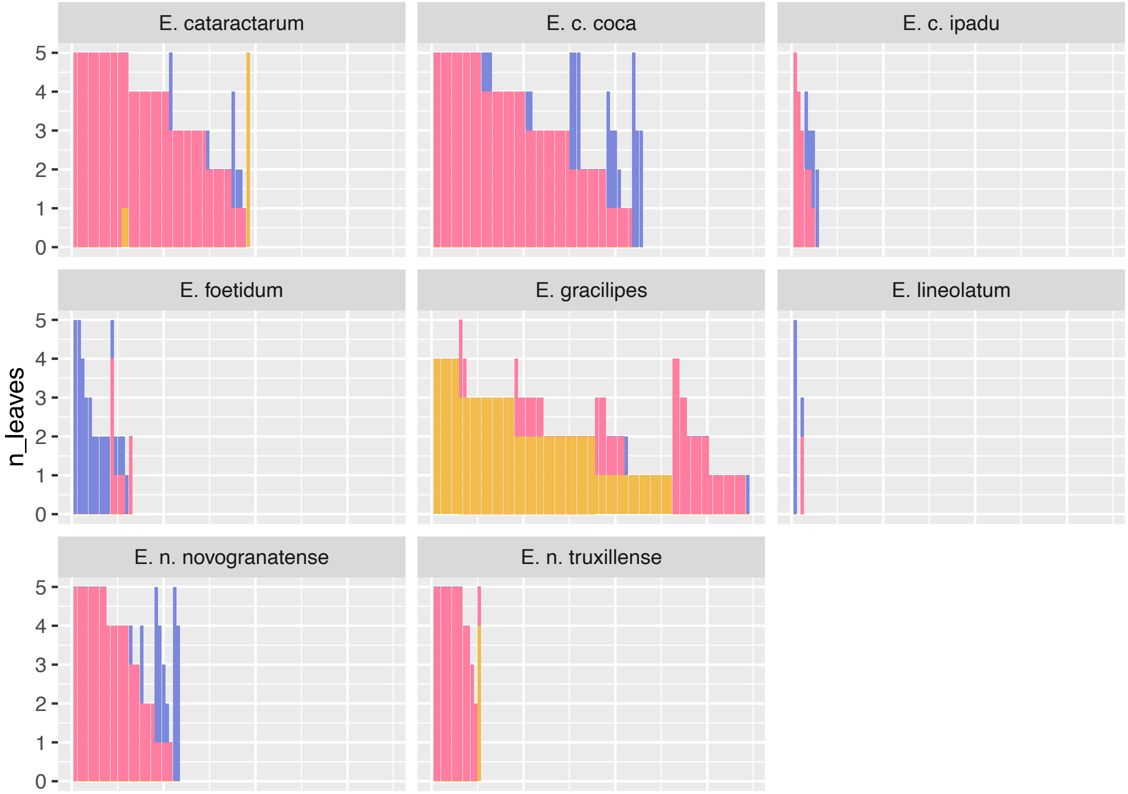


**Figure S8.** Bar charts representing the number of leaves sampled, and their individual cluster assignments based on GMM of EFA data (principal dataset of 835 leaves), and grouped by pre-defined taxonomic identification.

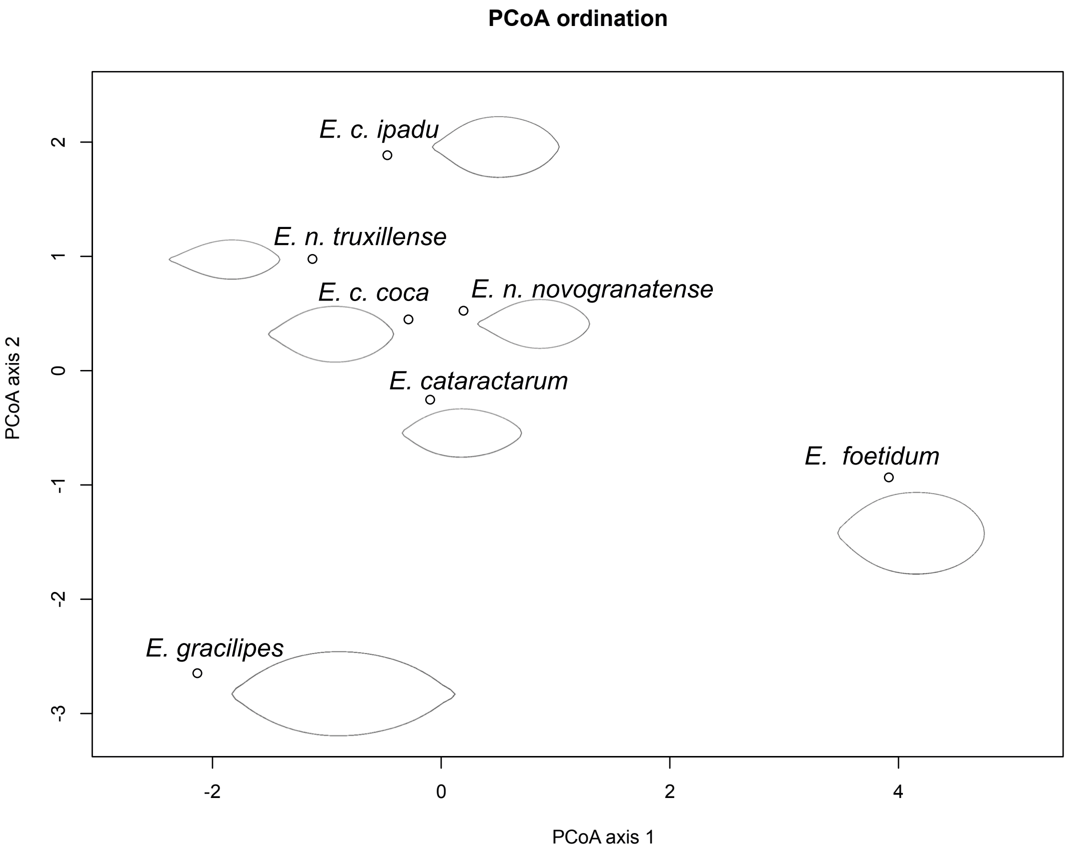


**Figure S9.** Principal Coordinates Analysis (PCoA) based on EFA data (principal dataset of 835 leaves) and representing centroids of recognised taxa.

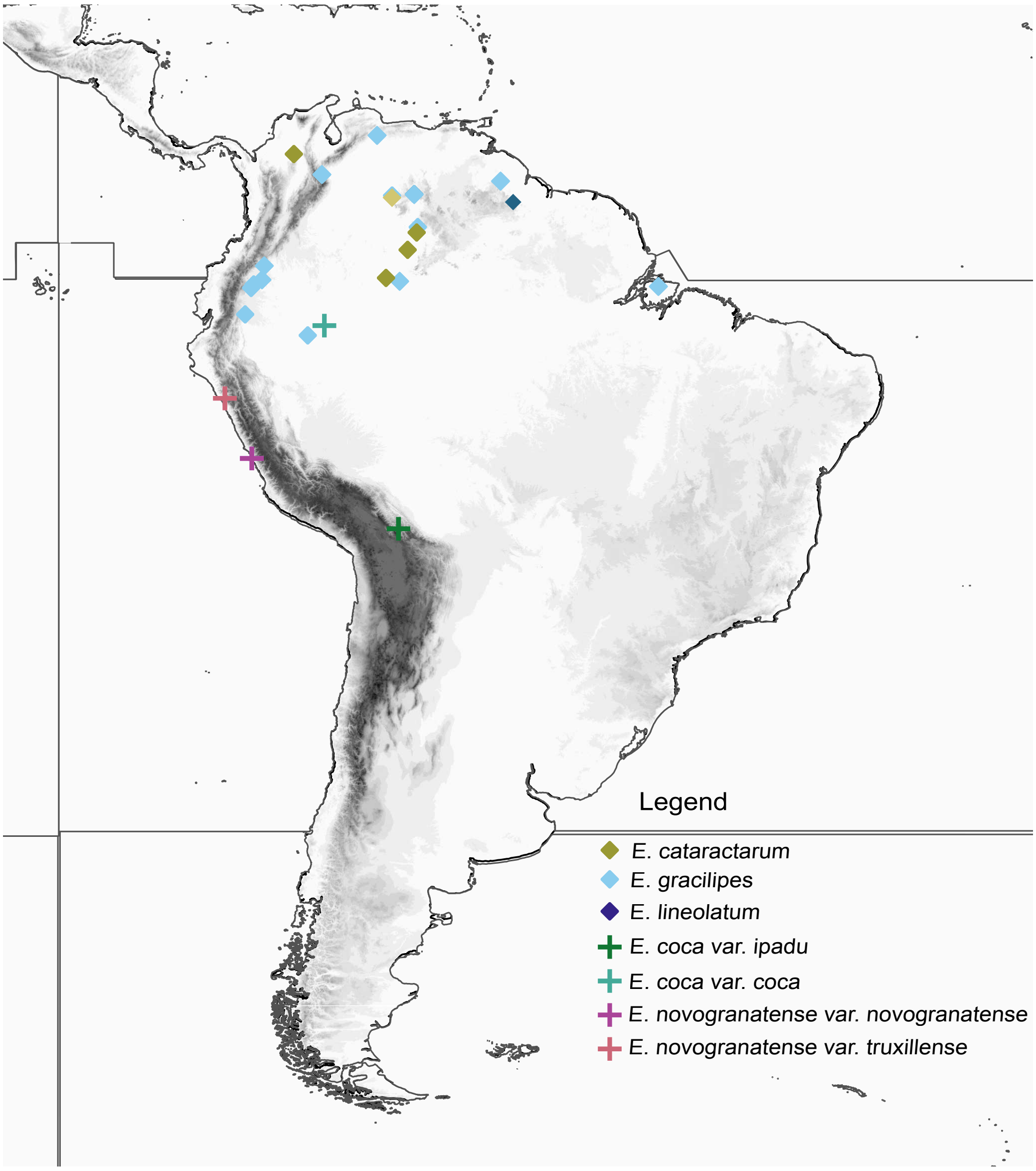


**Figure S10.** Geographical origins of *Erythroxylum* samples used in phylogenomic work (17 generated, 8 datamined – see Tables S2,3).

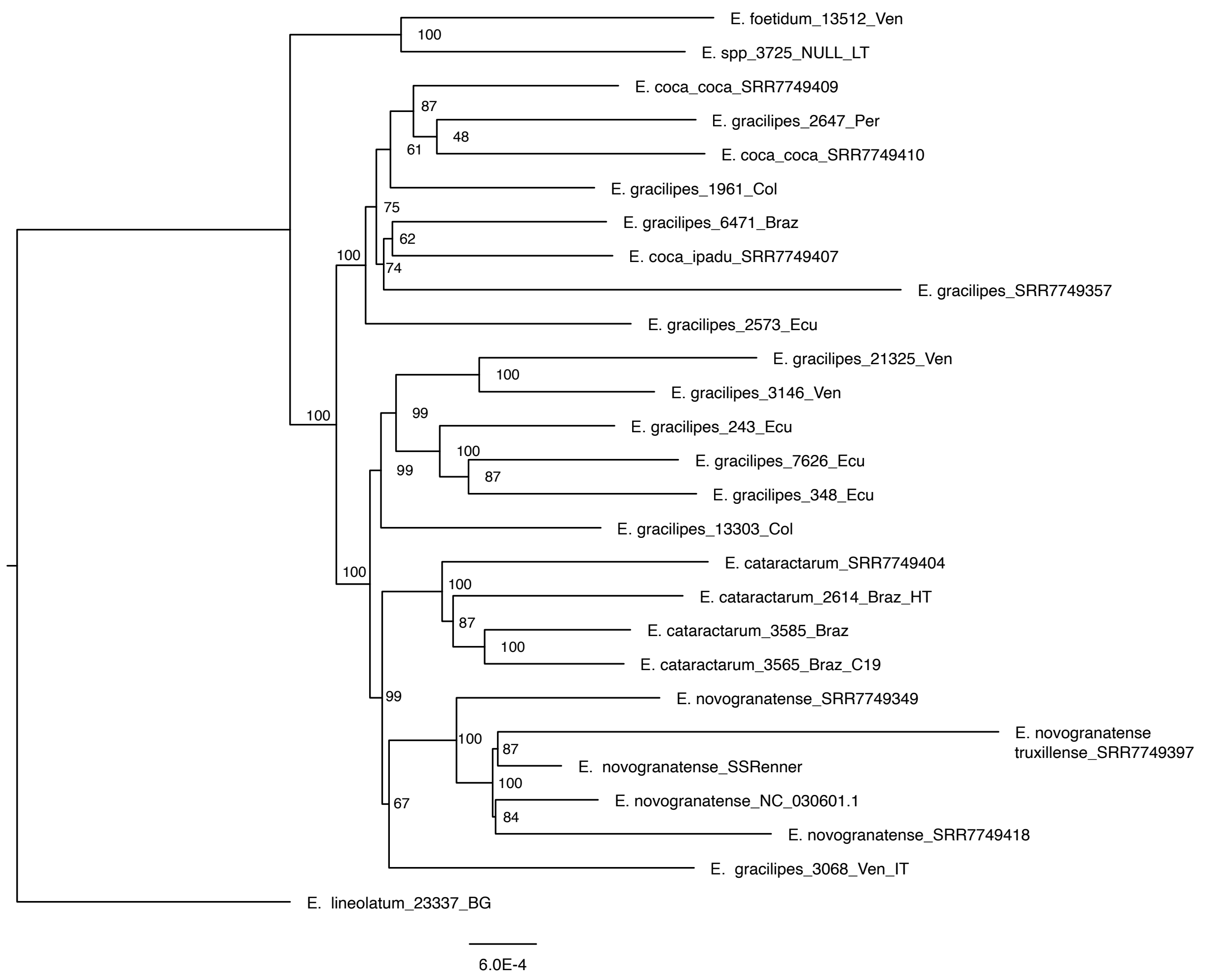


**Figure S11.** Maximum likelihood (ML) tree reconstructed from plastomes sequenced in 27 samples of *Erythroxylum,* with bootstrap support after 500 iterations indicated.

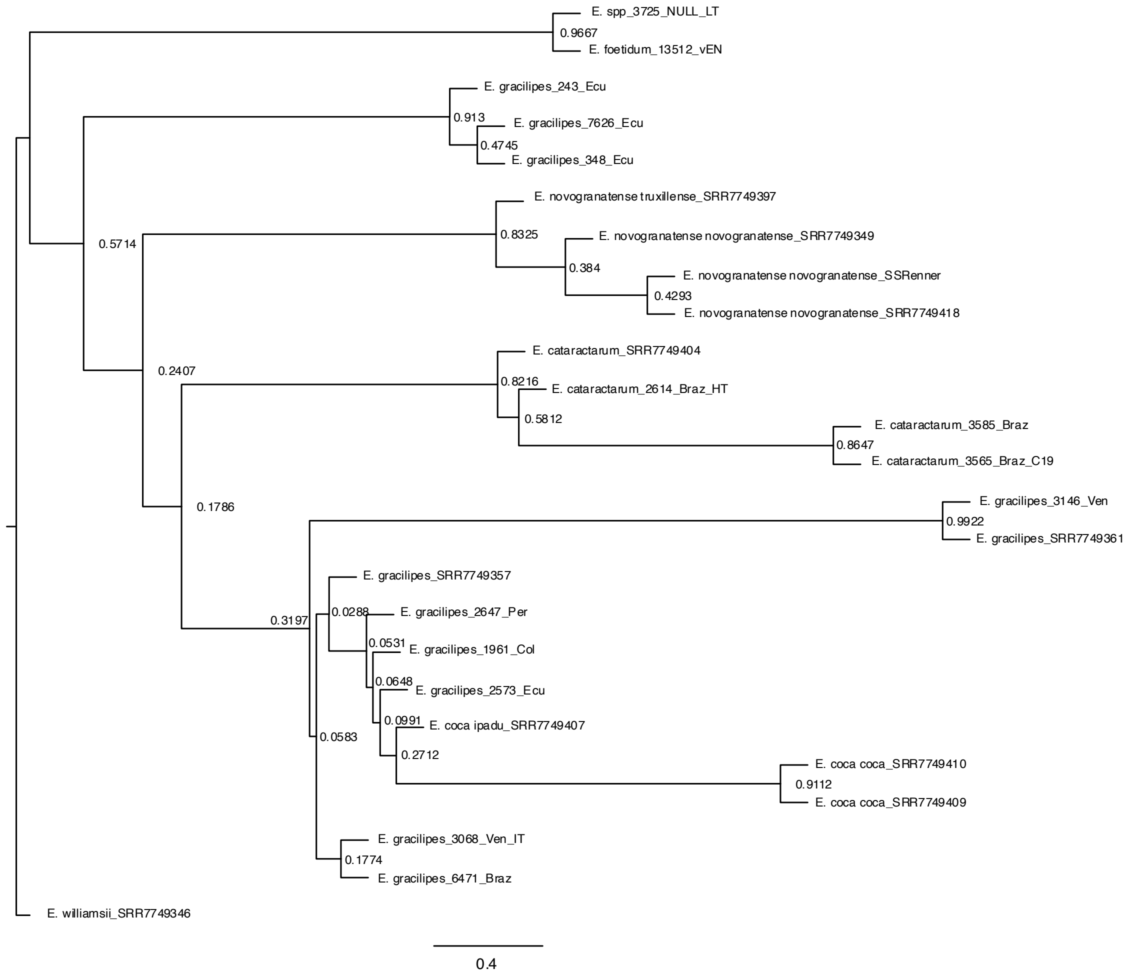


**Figure S12a.** ASTRAL summary species tree topology derived from 326 maximum likelihood (ML) phylogenies built from nuclear genes from 25 samples of *Erythroxylum*, with support indicated at each node as gene concordance factors (gCF).

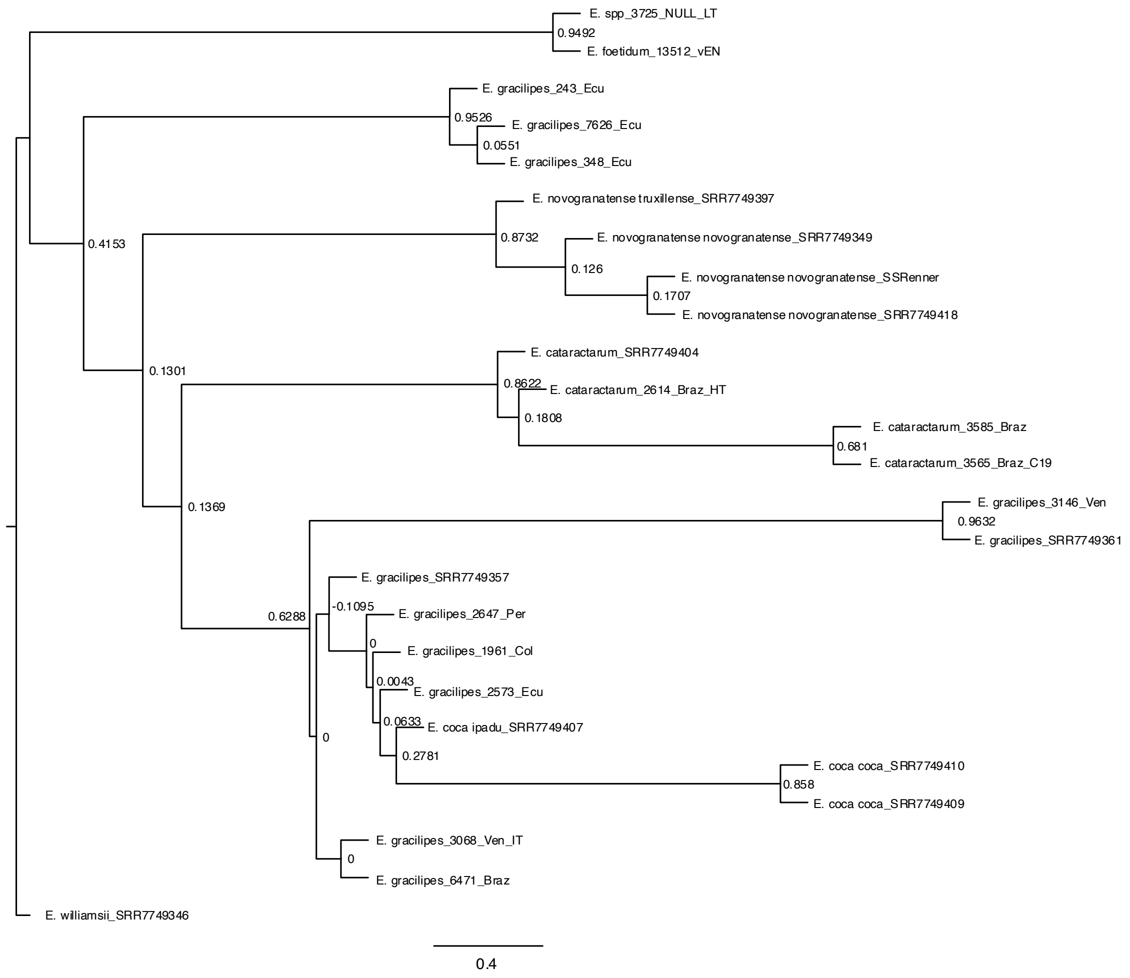


**Figure S12b.** ASTRAL summary species tree topology derived from 326 maximum likelihood (ML) phylogenies built from nuclear genes from 25 samples of *Erythroxylum*, with support indicated at each node as internode certainty (IC).

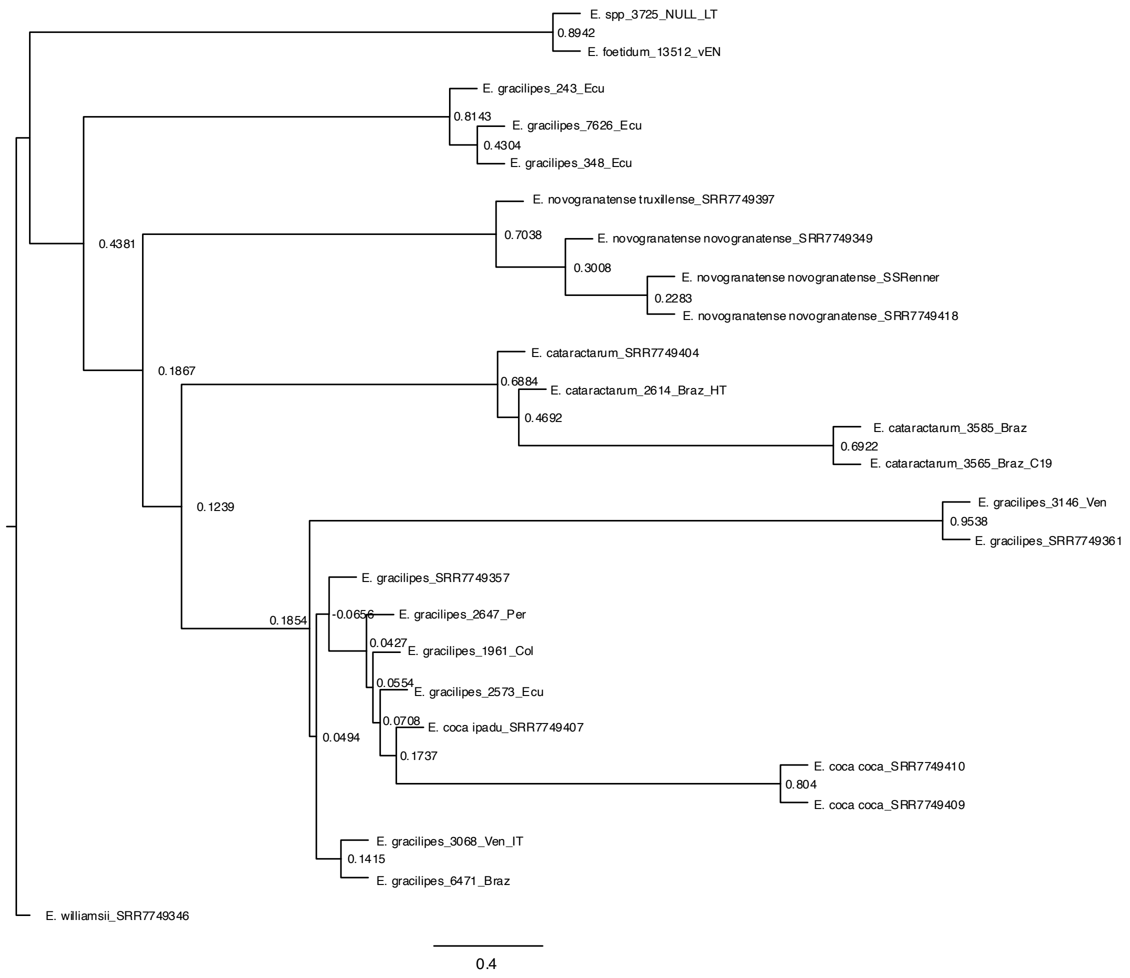


**Figure S12c.** ASTRAL summary species tree topology derived from 326 maximum likelihood (ML) phylogenies built from nuclear genes from 25 samples of *Erythroxylum*, with support indicated at each node as internode certainty ‘all’ (ICA).

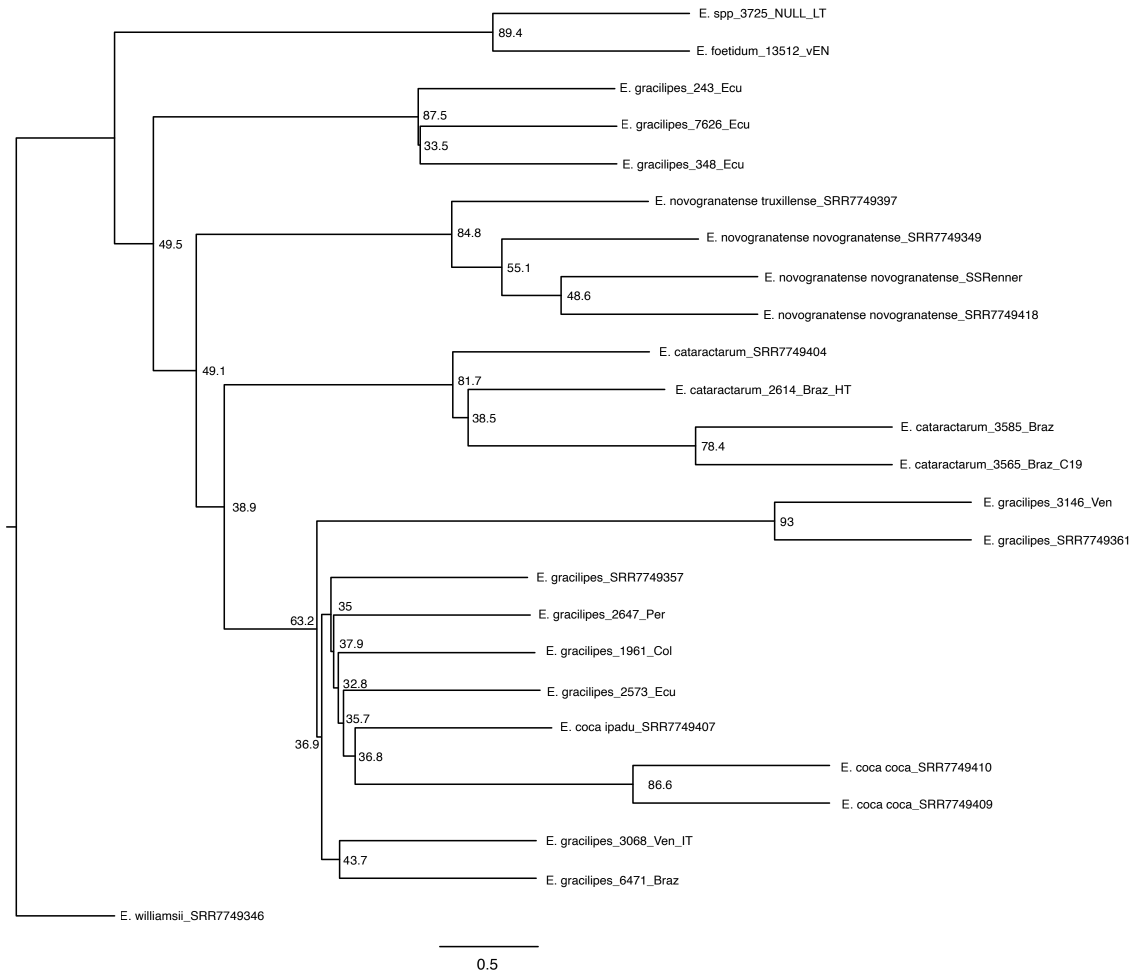


**Figure S12d.** ASTRAL summary species tree topology derived from 326 maximum likelihood (ML) phylogenies built from nuclear genes from 25 samples of *Erythroxylum*, with support indicated at each node as site concordance factors (sCF).

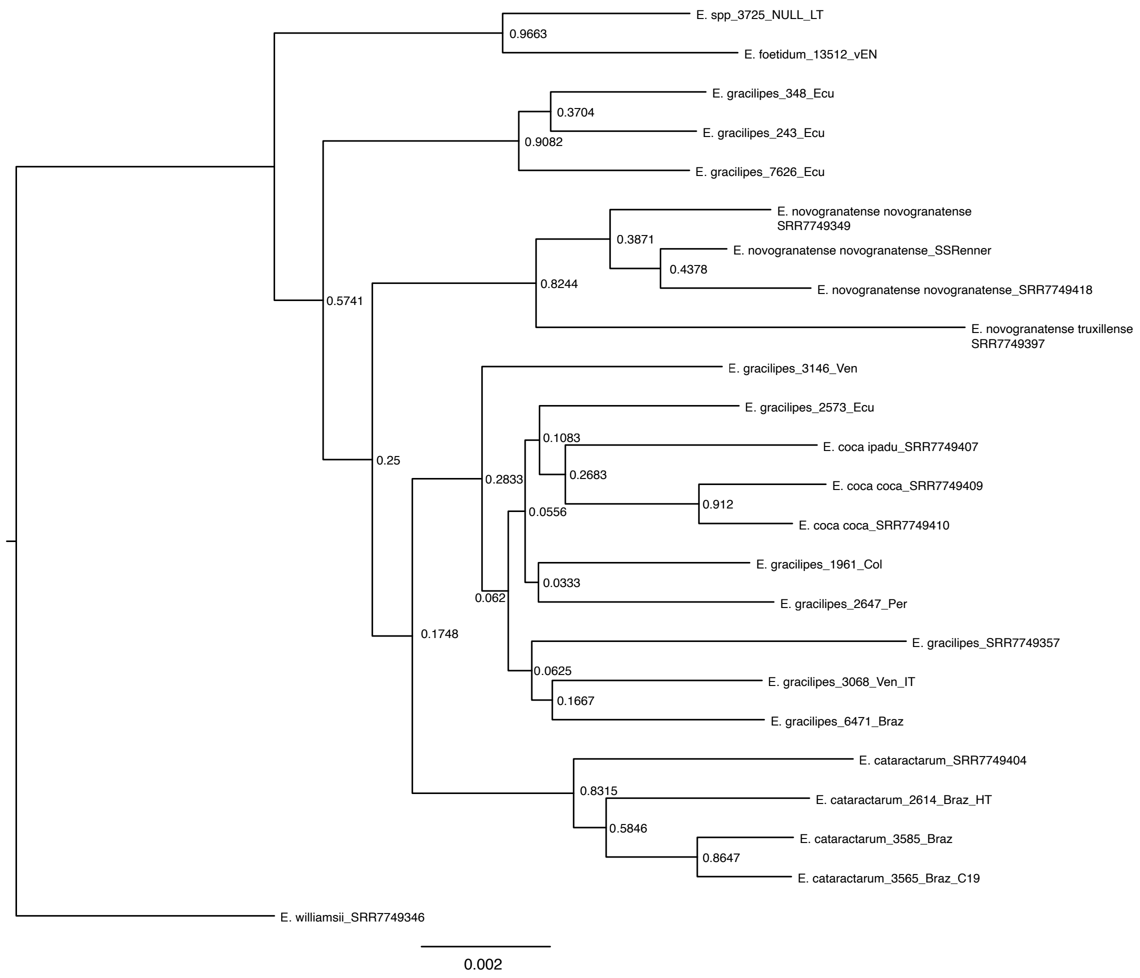


**Figure S13a.** RAxML summary species tree topology derived from a maximum likelihood (ML) phylogeny built from the concatenation of 326 nuclear gene alignments from 25 samples of *Erythroxylum*, with support indicated at each node as gene concordance factors (gCF).

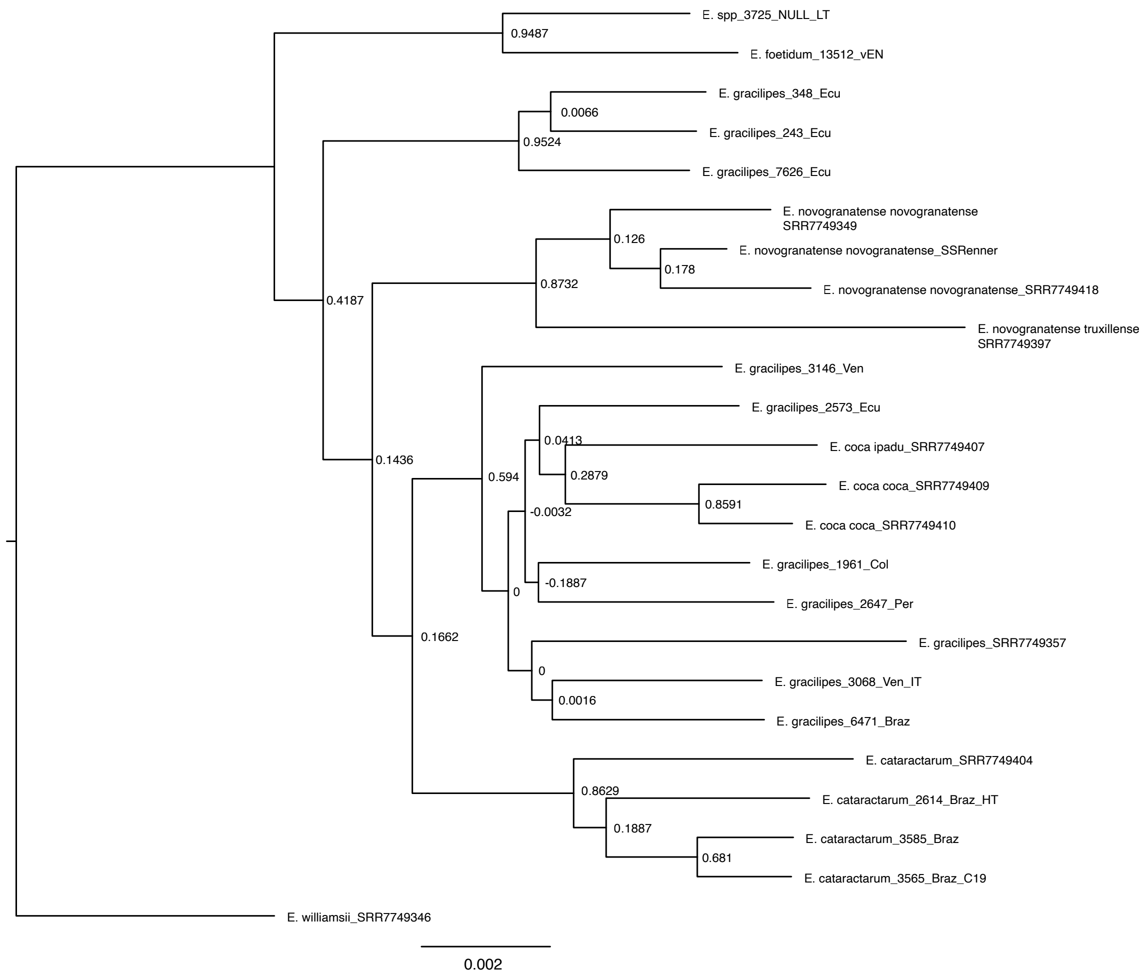


**Figure S13b.** RAxML summary species tree topology derived from a maximum likelihood (ML) phylogeny built from the concatenation of 326 nuclear gene alignments from 25 samples of *Erythroxylum*, with support indicated at each node as internode certainty (IC).

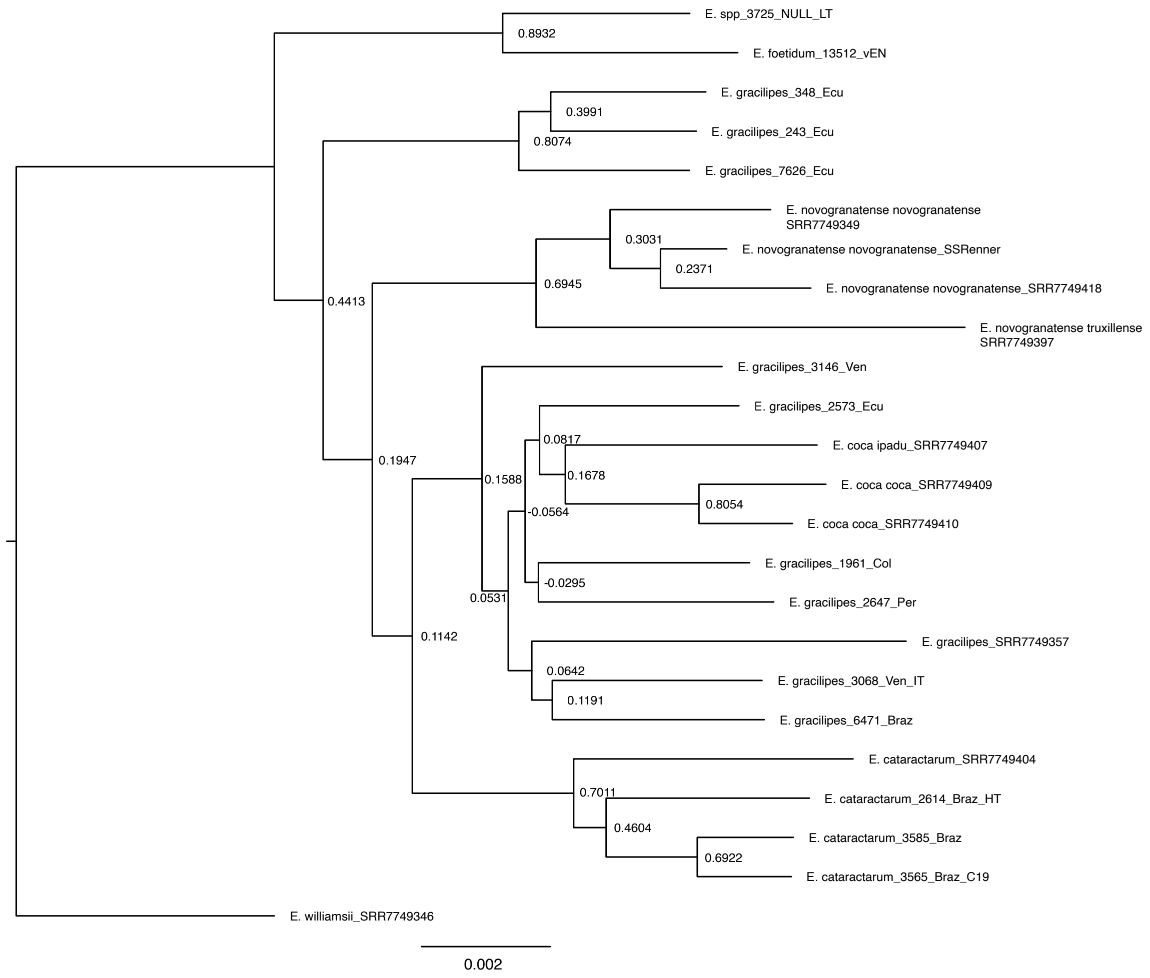


**Figure S13c.** RAxML summary species tree topology derived from a maximum likelihood (ML) phylogeny built from the concatenation of 326 nuclear gene alignments from 25 samples of *Erythroxylum*, with support indicated at each node as internode certainty 'all' (ICA).

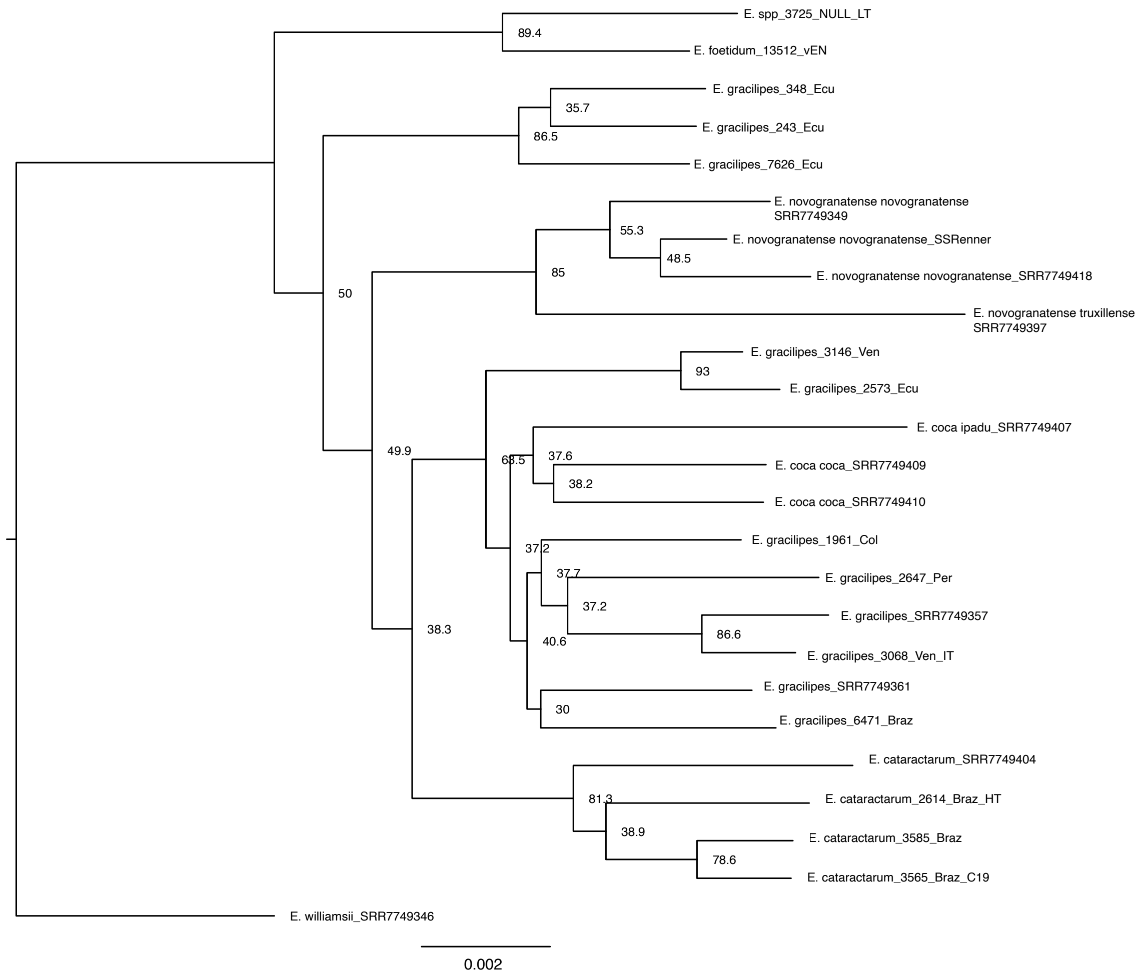


**Figure S13d.** RAxML summary species tree topology derived from a maximum likelihood (ML) phylogeny built from the concatenation of 326 nuclear gene alignments from 25 samples of *Erythroxylum*, with support indicated at each node as site concordance factors (sCF).

#### Phylogenetic network with hybrid events

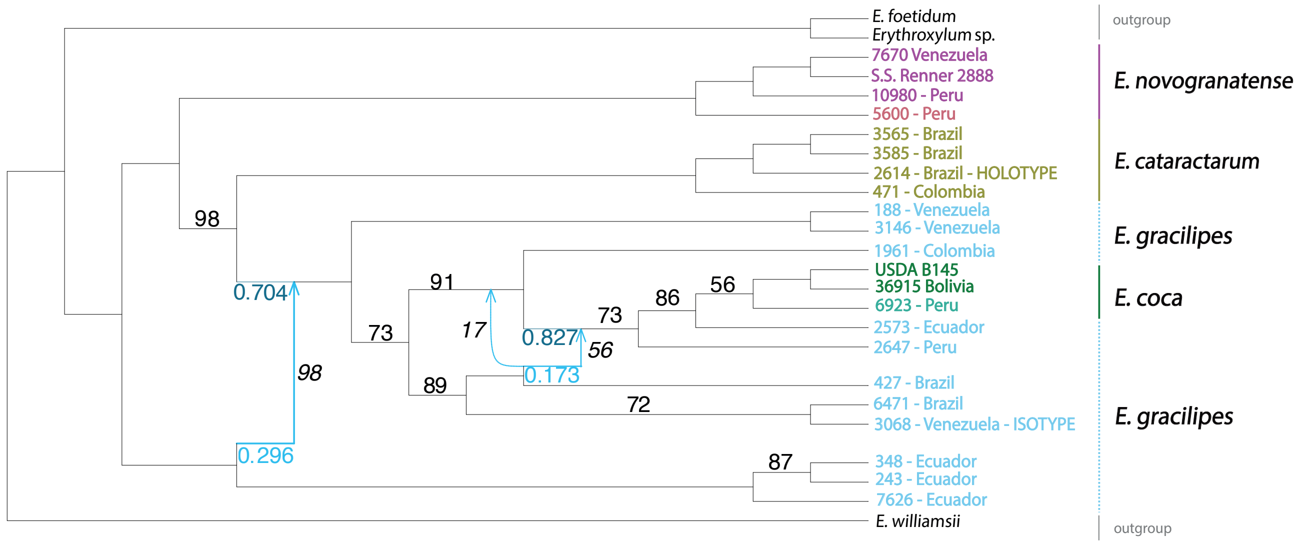


**Figure S14.** Phylogenetic network with two waves of gene flow ('hybrid events'). Numbers on branches represent bootstrap values lower than 100. Numbers in italics are bootstrap values for the hybridisations. The more basal hybridisation event was recovered in 98 of the bootstrap replicates, whereas the second one was recovered in 56 replicates. An alternative recipient position for that hybridisation is shown as a dotted line, which was recovered in 17 of the replicates. The direction of gene flow in the first hybridisation corresponds to the same bootstrap value and is therefore robust. The orientation of gene flow from the clade sister to *E. gracilipes* 427 (Brazil) has a bootstrap value of 74, i.e., the sum of bootstrap values for the minor edge in the optimal and alternative attachment places and is therefore also considered robust although highlighting some uncertainty. The dotted line represents an alternative position for the second hybridization event. Inheritance proportion values are presented for the optimal network. Coloured numbers represent the inheritance proportions for the major (dark blue) and minor (light blue) edges, depicting their point estimates. The credible intervals for the major and minor edges in the first hybridisation event were 0.624--0.796 and 0.204--0.374 respectively, whereas the intervals for the second hybridisation were 0.672--0.896 and 0.104--0.328.

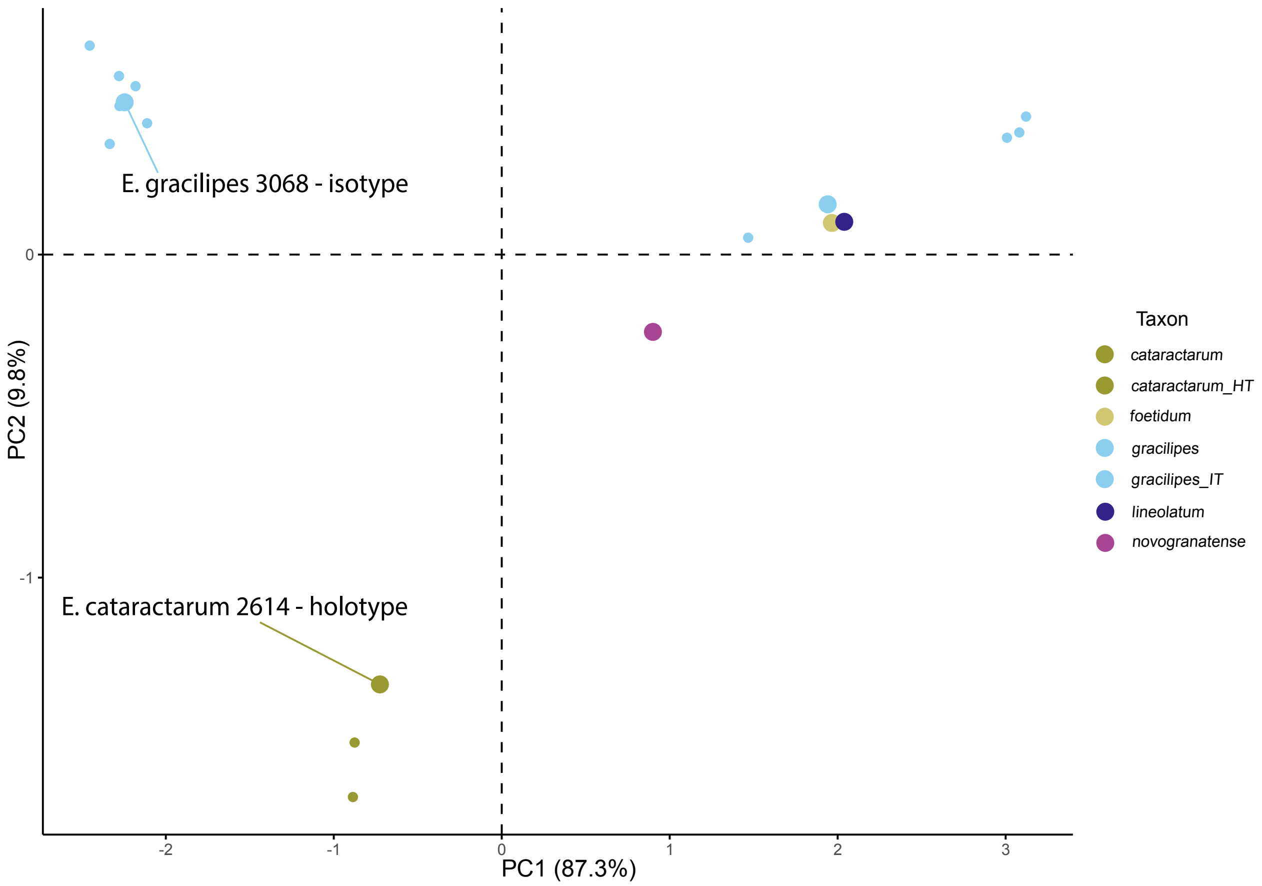


**Figure S15.** Nuclear genomic PCA based on 13,249 genotype likelihoods (GLs) computed from sequenced accessions of 20 *Erythroxylum* samples, predominantly sampled from the herbarium at RBG Kew (K).

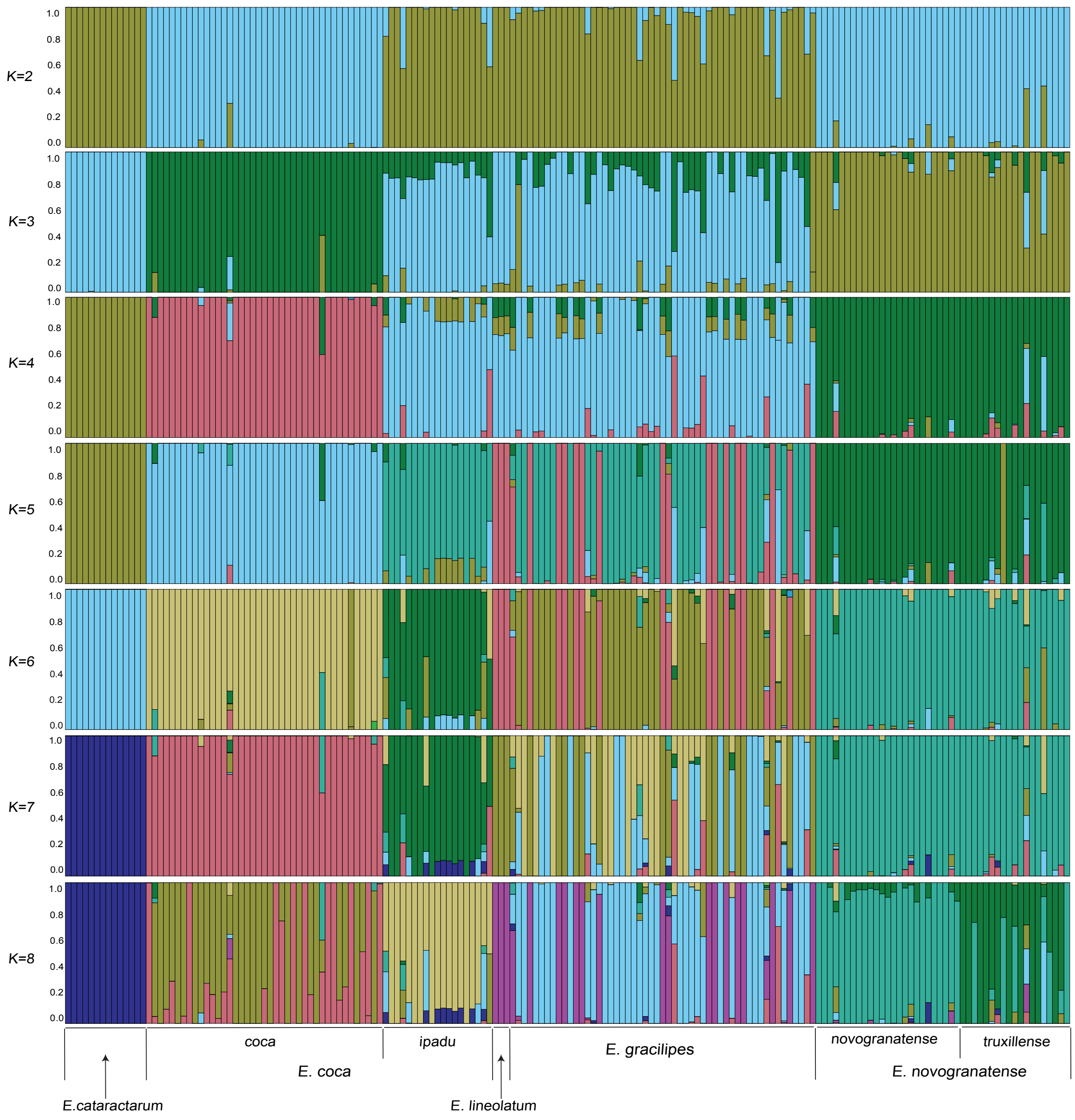


**Figure S16.** NGSadmix population structure inference showing per-individual ancestry of sequenced and datamined accessions of 173 *Erythroxylum* specimens at hypothetical values of *K* ranging from 2 to 10.

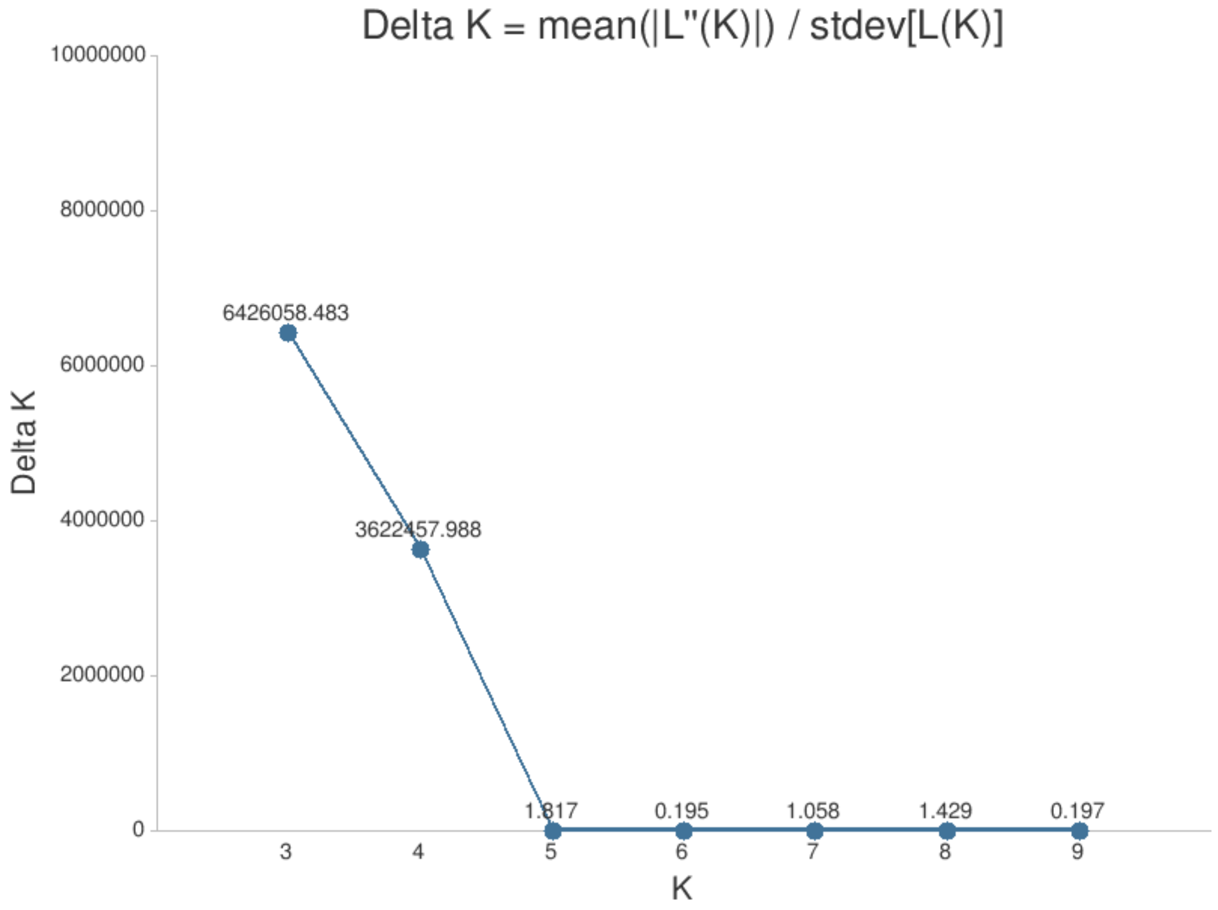


**Figure S17.** Theoretical best values of *K* inferred from output of NGSadmix population structure analysis, estimated using the Evanno method and implemented in CLUMPAK.

**Table S1. Taxon information and source herbarium for all specimens used in morphometric analyses. Herbarium acronyms are from Index Herbariorum (https://sweetgum.nybg.org/science/ih/).**

| Taxon (from label) | Catalogue Number | Collector | Collector number | Country of Collection | Herbarium | Taxon (post-verification) |
|---|---|---|---|---|---|---|
| *E. cataractarum* | BR0000006997533 | R. Spruce | 2614 | Brazil | BR | *E. cataractarum* |
| *E. cataractarum* | COL000373549 | J. Cuatrecasas | 13277 | Colombia | COL | *E. cataractarum* |
| *E. cataractarum* | V0055630F | R. Spruce | 2614 | Brazil | F | *E. cataractarum* |
| *E. cataractarum* | V0268796F | F. Guanchez | 3091 | Venezuela | F | *E. cataractarum* |
| *E. cataractarum* | V0268797F | E. Melgueiro | 136 | Venezuela | F | *E. cataractarum* |
| *E. cataractarum* | V0268798F | R. S. Cowan | 32069 | Venezuela | F | *E. cataractarum* |
| *E. cataractarum* | V0268799F | F. Guanchez | 3182 | Venezuela | F | *E. cataractarum* |
| *E. cataractarum* | V0268801F | F. Guanchez | 3093 | Venezuela | F | *E. cataractarum* |
| *E. cataractarum* | V0268802F | F. Guanchez | 3092 | Venezuela | F | *E. cataractarum* |
| *E. cataractarum* | V0268804F | G. Davidse | 15626 | Venezuela | F | *E. cataractarum* |
| *E. cataractarum* | V0268806F | J. Zarucchi | 1422 | Colombia | F | *E. cataractarum* |
| *E. cataractarum* | V0268807F | F. Guanchez | 2938 | Venezuela | F | *E. cataractarum* |
| *E. cataractarum* | V0268809F | F. Guanchez | 3090 | Venezuela | F | *E. cataractarum* |
| *E. cataractarum* | V0268810F | G. Davidse | 1413 | Venezuela | F | *E. cataractarum* |
| *E. cataractarum* | V0268811F | R. Callejas | 4371 | Colombia | F | *E. cataractarum* |
| *E. cataractarum* | V0268812F | F. Delascioo | 11189 | Venezuela | F | *E. cataractarum* |
| *E. cataractarum* | V0268813F | B. Stergios | 8231 | Colombia | F | *E. cataractarum* |
| *E. cataractarum* | V0268816F | G. Davidse | 15975 | Venezuela | F | *E. cataractarum* |
| *E. cataractarum* | V0268817F | J. Zarucchi | 1268 | Colombia | F | *E. cataractarum* |
| *E. cataractarum* | V0268818F | G. Davidse | 15814 | Venezuela | F | *E. cataractarum* |
| *E. cataractarum* | V0268819F | J. Zarucchi | 1383 | Colombia | F | *E. cataractarum* |
| *E. cataractarum* | V0268823F | W. Davis | 151 | Colombia | F | *E. cataractarum* |
| *E. cataractarum* | V0272206F | R. Liesner | 11892 | Venezuela | F | *E. cataractarum* |
| *E. cataractarum* | V0471772F | D.M. White | 553 | Colombia | F | *E. cataractarum* |
| *E. cataractarum* | V0471774F | D.M. White | 615 | Colombia | F | *E. cataractarum* |
| *E. cataractarum* | V0471775F | D.M. White | 551 | Colombia | F | *E. cataractarum* |
| *E. cataractarum* | V0471777F | D.M. White | 558 | Colombia | F | *E. cataractarum* |
| *E. cataractarum* | K000426845 | R. Spruce | 2614 | Brazil | K | *E. cataractarum* |
| *E. cataractarum* | K001201221 | R. Spruce | 3585 | Brazil | K | *E. cataractarum* |
| *E. cataractarum* | s.n. | J. Zarucchi | 1383 | Colombia | K | *E. cataractarum* |
| *E. cataractarum* | s.n. | J. Zarucchi | 1422 | Colombia | K | *E. cataractarum* |
| *E. cataractarum* | s.n. | R. Spruce | 3565 | Brazil | K | *E. cataractarum* |
| *E. cataractarum* | s.n. | T. Plowman | 4265 | Colombia | K | *E. cataractarum* |
| *E. cataractarum* | s.n. | W. Davis | 151 | Colombia | K | *E. cataractarum* |
| *E. cataractarum* | L3784047 | G. Davidse | 15814 | Venezuela | L | *E. cataractarum* |
| *E. cataractarum* | P05483127 | G. Davidse | 14313 | Venezuela | P | *E. cataractarum* |
| *E. cataractarum* | U1284837 | R. Schultes | 16196 | Venezuela | U | *E. cataractarum* |
| *E. cataractarum* | 288464 | J. Zarucchi | 1422 | Colombia | US | *E. cataractarum* |

| | | | | | | |
|---|---|---|---|---|---|---|
| *E. cataractarum* | 1773994 | J. Cuatrecasas | 4337 | Colombia | US | *E. cataractarum* |
| *E. cataractarum* | 2171745 | R. Schultes | 16194 | Venezuela | US | *E. cataractarum* |
| *E. cataractarum* | 2171746 | R. Schultes | 16196 | Venezuela | US | *E. cataractarum* |
| *E. cataractarum* | 2253071 | R.S. Cowan | 32069 | Venezuela | US | *E. cataractarum* |
| *E. cataractarum* | 2458022 | L. Uribe | 5183 | Colombia | US | *E. cataractarum* |
| *E. cataractarum* | 2969895 | J. Zarucchi | 1383 | Colombia | US | *E. cataractarum* |
| *E. cataractarum* | 2969896 | J. Zarucchi | 1422 | Colombia | US | *E. cataractarum* |
| *E. cataractarum* | 3108061 | F. Delascioo | 11189 | Venezuela | US | *E. cataractarum* |
| *E. coca* var. *coca* | B100157443 | P. Goodell | 75-50 | Peru | B | *E. coca* var. *coca* |
| *E. coca* var. *coca* | BISH1010745 | T. Flynn | 476 | USA (Hawaii) | BISH | *E. n.* var. *novogranatense* |
| *E. coca* var. *coca* | BISH1010746 | F.R.F. | 35255 | Palau | BISH | *E. n.* var. *novogranatense* |
| *E. coca* var. *coca* | BISH1010748 | L. Williams | 3551 | Peru | BISH | *E. coca* var. *coca* |
| *E. coca* var. *coca* | BR0000014613401 | A. Flamigni | 13/A | Congo, Democratic Republic | BR | *E. coca* var. *coca* |
| *E. coca* var. *coca* | BR0000014613432 | M. Laurent | s.n. | Congo, Democratic Republic | BR | *E. coca* var. *coca* |
| *E. coca* var. *coca* | BR0000014613456 | M. Laurent | s.n. | Belgium | BR | *E. n.* var. *novogranatense* |
| *E. coca* var. *coca* | BR0000014613463 | M. Laurent | s.n. | Belgium | BR | *E. n.* var. *novogranatense* |
| *E. coca* var. *coca* | BR0000014613494 | F. Vermoesen | 2080 | Congo, Democratic Republic | BR | *E. coca* var. *coca* |
| *E. coca* var. *coca* | BR0000014613500 | F. Vermoesen | 2081 | Congo, Democratic Republic | BR | *E. coca* var. *coca* |
| *E. coca* var. *coca* | BR0000014613548 | A.L.B. Eala | 1946 | Congo, Democratic Republic | BR | *E. coca* var. *coca* |
| *E. coca* var. *coca* | BR0000014613555 | A. Corbisier-Baland | 1946 | Congo, Democratic Republic | BR | *E. coca* var. *coca* |
| *E. coca* var. *coca* | BR0000014613579 | É. Laurent | s.n. | indet. | BR | *E. coca* var. *coca* |
| *E. coca* var. *coca* | BR0000014613593 | A. Corbisier-Baland | 1058 | Congo, Democratic Republic | BR | *E. coca* var. *coca* |
| *E. coca* var. *coca* | BR0000014613609 | A. Corbisier-Baland | 1059 | Congo, Democratic Republic | BR | *E. coca* var. *coca* |
| *E. coca* var. *coca* | BR0000014613647 | A. Corbisier-Baland | 1941 | Congo, Democratic Republic | BR | *E. coca* var. *coca* |
| *E. coca* var. *coca* | NCY016238 | A. Bour | s.n. | France | CJBN | *E. coca* var. *coca* |
| *E. coca* var. *coca* | ESA004936 | W.R., Accorsi | s.n. | Brazil | ESA | *E. coca* var. *coca* |
| *E. coca* var. *coca* | V0321195F | D.M. White | 490 | Peru | F | *E. coca* var. *coca* |
| *E. coca* var. *coca* | IAN043388 | A. Ducke | s.n. | Brazil | IAN | *E. coca* var. *coca* |
| *E. coca* var. *coca* | IAN050048 | J.M. Pires | 1870 | Brazil | IAN | *E. coca* var. *coca* |
| *E. coca* var. *coca* | K000700870 | without collector | s.n. | Sri Lanka | K | *E. coca* var. *coca* |

| | | | | | | |
|---|---|---|---|---|---|---|
| *E. coca* var. *coca* | L2120521 | F. Sandberg | 1959-466 | Peru | L | *E. coca* var. *coca* |
| *E. coca* var. *coca* | L2120522 | E. Ule | 5039 | Brazil | L | *E. coca* var. *coca* |
| *E. coca* var. *coca* | L2120606 | without collector | s.n. | indet. | L | *E. coca* var. *coca* |
| *E. coca* var. *coca* | L2120611 | without collector | s.n. | indet. | L | *E. coca* var. *coca* |
| *E. coca* var. *coca* | MA-01-00812602 | without collector | s.n. | indet. | MA | *E. coca* var. *coca* |
| *E. coca* var. *coca* | MA-01-00812603 | without collector | s.n. | Peru | MA | *E. coca* var. *coca* |
| *E. coca* var. *coca* | MA-01-00812604 | without collector | s.n. | Peru | MA | *E. coca* var. *coca* |
| *E. coca* var. *coca* | MA-01-00817422 | without collector | s.n. | indet. | MA | *E. coca* var. *coca* |
| *E. coca* var. *coca* | MA-01-00818690 | without collector | s.n. | indet. | MA | *E. coca* var. *coca* |
| *E. coca* var. *coca* | MA-01-00818691 | without collector | s.n. | indet. | MA | *E. coca* var. *coca* |
| *E. coca* var. *coca* | MA-01-00818692 | without collector | s.n. | indet. | MA | *E. coca* var. *coca* |
| *E. coca* var. *coca* | MA-01-00818693 | without collector | s.n. | indet. | MA | *E. coca* var. *coca* |
| *E. coca* var. *coca* | MA-01-00818694 | without collector | s.n. | indet. | MA | *E. coca* var. *coca* |
| *E. coca* var. *coca* | MA-01-00818696 | without collector | s.n. | indet. | MA | *E. coca* var. *coca* |
| *E. coca* var. *coca* | MA-01-00818697 | without collector | s.n. | indet. | MA | *E. coca* var. *coca* |
| *E. coca* var. *coca* | MA-01-00818698 | without collector | s.n. | indet. | MA | *E. coca* var. *coca* |
| *E. coca* var. *coca* | MA-01-00818700 | without collector | s.n. | indet. | MA | *E. coca* var. *coca* |
| *E. coca* var. *coca* | MA-01-00818701 | without collector | s.n. | indet. | MA | *E. coca* var. *coca* |
| *E. coca* var. *coca* | MBM279737 | R. Mello-Silva | 2130 | Bolivia | MBM | *E. coca* var. *coca* |
| *E. coca* var. *coca* | MG030282 | N. Pereira | 30282 | Brazil | MG | *E. coca* var. *coca* |
| *E. coca* var. *coca* | MG084033 | J. Jangoux | 1221 | Brazil | MG | *E. coca* var. *coca* |
| *E. coca* var. *coca* | MG100902 | B. Nelson | 422 | Brazil | MG | *E. coca* var. *coca* |
| *E. coca* var. *coca* | P05480975 | without collector | 1700 | Cameroon | P | *E. coca* var. *coca* |
| *E. coca* var. *coca* | P05480981 | A. Corbisier-Baland | 1058 | Congo, Democratic Republic | P | *E. coca* var. *coca* |
| *E. coca* var. *coca* | P05483609 | C.F. Baker | 133 | Sri Lanka | P | *E. coca* var. *coca* |
| *E. coca* var. *coca* | P05485753 | C. Sastre | 2459 | Colombia | P | *E. coca* var. *coca* |
| *E. coca* var. *coca* | P05486617 | without collector | s.n. | indet. | P | *E. coca* var. *coca* |
| *E. coca* var. *coca* | P05486619 | J.S. Vigo | 10018 | Peru | P | *E. coca* var. *coca* |
| *E. coca* var. *coca* | P05486775 | without collector | s.n. | indet. | P | *E. n.* var. *novogranatense* |
| *E. coca* var. *coca* | P05576214 | T. Plowman | 8029 | USA | P | *E. coca* var. *coca* |
| *E. coca* var. *coca* | P05576223 | without collector | s.n. | Peru | P | *E. coca* var. *coca* |
| *E. coca* var. *coca* | P05576231 | W. Lechler | 2220 | indet. | P | *E. coca* var. *coca* |

| | | | | | | |
|---|---|---|---|---|---|---|
| *E. coca* var. *coca* | P05576237 | A. Glaziou | 14554 | Brazil | P | *E. coca* var. *coca* |
| *E. coca* var. *coca* | P05576239 | C.F. Baker | 70 | Brazil | P | *E. coca* var. *coca* |
| *E. coca* var. *coca* | P05576246 | M. Weddell | 4265 | Bolivia | P | *E. coca* var. *coca* |
| *E. coca* var. *coca* | P05600998 | without collector | IX | indet. | P | *E. n.* var. *novogranatense* |
| *E. coca* var. *coca* | 2794875 | L.Ma.M. Carreira | 203 | Brazil | MO | *E. coca* var. *coca* |
| *E. coca* var. *coca* | 6606818 | M. Monigatti | 79 | Peru | MO | *E. coca* var. *coca* |
| *E. coca* var. *coca* | NCSC00014110 | E. Machado | 1105 | Peru | NCSC | *E. coca* var. *coca* |
| *E. coca* var. *coca* | NCSC00014117 | A. Marchena | 1523 | Peru | NCSC | *E. coca* var. *coca* |
| *E. coca* var. *coca* | PH00026753 | M. Bang | 268 | Bolivia | PH | *E. coca* var. *coca* |
| *E. coca* var. *coca* | RB00078682 | R. Mello-Silva | 2130 | Bolivia | RB | *E. coca* var. *coca* |
| *E. coca* var. *coca* | RB00078712 | P. Occhioni | 4813 | indet. | RB | *E. coca* var. *coca* |
| *E. coca* var. *coca* | RB00078761 | J. Schunke | 10018 | Peru | RB | *E. coca* var. *coca* |
| *E. coca* var. *coca* | RB00078783 | T. Plowman | 5833 | Peru | RB | *E. coca* var. *coca* |
| *E. coca* var. *coca* | RB00078803 | L.Ma.M. Carreira | 203 | Brazil | RB | *E. coca* var. *coca* |
| *E. coca* var. *coca* | RB00783565 | V.F. Kinupp | 3638 | Brazil | RB | *E. coca* var. *coca* |
| *E. coca* var. *coca* | SPF00156843 | R. Mello-Silva | 2130 | Bolivia | SPF | *E. coca* var. *coca* |
| *E. coca* var. *coca* | U1270510 | J.M. Pires | 51919 | Brazil | U | *E. coca* var. *coca* |
| *E. coca* var. *coca* | U1284615 | M. Buysman | 6 | Grenada | U | *E. coca* var. *coca* |
| *E. coca* var. *coca* | U1284756 | Y. Mexia | 7795 | Bolivia | U | *E. coca* var. *coca* |
| *E. coca* var. *coca* | U1284757 | T. Plowman | 7540 | Peru | U | *E. coca* var. *coca* |
| *E. coca* var. *coca* | U1284778 | W. Davis | 13 | Colombia | U | *E. coca* var. *coca* |
| *E. coca* var. *coca* | U1284779 | G. Meyer | 341 | Bolivia | U | *E. n.* var. *truxillense* |
| *E. coca* var. *coca* | U1284782 | A. Gentry | 15975 | Peru | U | *E. coca* var. *coca* |
| *E. coca* var. *coca* | U1603902 | without collector | 68-714 | indet. | U | *E. n.* var. *novogranatense* |
| *E. coca* var. *coca* | H1235777 | F. Ganders | 79-6 | Canada | UAM | *E. coca* var. *coca* |
| *E. coca* var. *coca* | V172500 | F. Ganders | 79-7 | Canada | UBC | *E. coca* var. *coca* |
| *E. coca* var. *coca* | V172508 | T.A. Johns | 330 ab | Peru | UBC | *E. coca* var. *coca* |
| *E. coca* var. *coca* | V193191 | T. Plowman | 5975 | Peru | UBC | *E. coca* var. *coca* |
| *E. coca* var. *coca* | 97511 | W. Gillis | 7558 | USA | USF | *E. coca* var. *coca* |
| *E. coca* var. *coca* | 170609 | J. Zarucchi | 1145 | Colombia | USF | *E. coca* var. *coca* |
| *E. coca* var. *coca* | 190941 | R. Vásquez | 5793 | Peru | USF | *E. coca* var. *coca* |
| *E. coca* var. *coca* | UTC00203717 | C. McMullen | 697 | Bolivia | UTC | *E. coca* var. *coca* |
| *E. coca* var. *coca* | W0059814 | Poeppig | s.n. | Brazil | W | *E. coca* var. *coca* |
| *E. coca* var. *ipadu* | COL000373556 | J. Cuatrecasas | 7012 | Colombia | COL | *E. coca* var. *ipadu* |
| *E. coca* var. *ipadu* | ECON00043856 | T. Plowman | 6663 | Peru | ECON | *E. coca* var. *ipadu* |
| *E. coca* var. *ipadu* | V0268821F | J. Ruíz | 1254 | Peru | F | *E. coca* var. *ipadu* |
| *E. coca* var. *ipadu* | GH00043857 | T. Plowman | 4463 | Peru | GH | *E. coca* var. *ipadu* |
| *E. coca* var. *ipadu* | IAN151984 | W. Davis | 20 | Colombia | IAN | *E. coca* var. *ipadu* |
| *E. coca* var. *ipadu* | K001202658 | Burchell | 9588 | Brazil | K | *E. coca* var. *ipadu* |
| *E. coca* var. *ipadu* | K001202659 | E. Ule | 5039 | Brazil | K | *E. coca* var. *ipadu* |

| | | | | | | |
|---|---|---|---|---|---|---|
| *E. coca* var. *ipadu* | K001202661 | M.A. Glaziou | 7537 | Brazil | K | *E. coca* var. *ipadu* |
| *E. coca* var. *ipadu* | K001202662 | G.T. Prance | 15572 | Brazil | K | *E. coca* var. *ipadu* |
| *E. coca* var. *ipadu* | K001202663 | T. Plowman | 8055 | Brazil | K | *E. coca* var. *ipadu* |
| *E. coca* var. *ipadu* | K001202664 | J.M. Pires | 51919 | Brazil | K | *E. coca* var. *ipadu* |
| *E. coca* var. *ipadu* | MG030222 | P. Cavalcante | 30222 | Brazil | MG | *E. coca* var. *ipadu* |
| *E. coca* var. *ipadu* | P05576204 | Jobert | m-9670 | Brazil | P | *E. coca* var. *ipadu* |
| *E. coca* var. *ipadu* | P05576206 | Jobert | s.n. | Brazil | P | *E. coca* var. *ipadu* |
| *E. coca* var. *ipadu* | P05576210 | E. Poisson | s.n. | Brazil | P | *E. coca* var. *ipadu* |
| *E. coca* var. *ipadu* | RB00078688 | unreadable | 7640 | Brazil | RB | *E. coca* var. *ipadu* |
| *E. coca* var. *ipadu* | RB00078702 | J. Jangoux | 1221 | Brazil | RB | *E. coca* var. *ipadu* |
| *E. coca* var. *ipadu* | RB00078806 | W. Rodrigues | 102931 | indet | RB | *E. coca* var. *ipadu* |
| *E. coca* var. *ipadu* | U1284784 | J. Torres | 167 | Peru | U | *E. coca* var. *ipadu* |
| *E. coca* var. *ipadu* | US00101253 | T. Plowman | 6663 | Peru | US | *E. coca* var. *ipadu* |
| *E. coca* var. *ipadu* | WAG1930897 | T. Plowman | 10934 | USA | WAG | *E. coca* var. *ipadu* |
| *E. coca* var. *ipadu* | WAG1930899 | C.L.M. van Eijnatten | 1849 | Benin | WAG | *E. coca* var. *ipadu* |
| *E. foetidum* | 1863255 | G. Davidse | 15280 | Venezuela | F | *E. foetidum* |
| *E. foetidum* | 1932921 | T. Plowman | 13510 | Venezuela | F | *E. foetidum* |
| *E. foetidum* | V0269780F | G. Davidse | 15593 | Venezuela | F | *E. foetidum* |
| *E. foetidum* | V0269781F | T. Plowman | 13731 | Venezuela | F | *E. foetidum* |
| *E. foetidum* | V0269784F | T. Plowman | 13731 | Venezuela | F | *E. foetidum* |
| *E. foetidum* | V0269789F | J. Steyermark | 122557 | Venezuela | F | *E. foetidum* |
| *E. foetidum* | V0269793F | G. Bunting | 7571 | Venezuela | F | *E. foetidum* |
| *E. foetidum* | V0269795F | G. Davidse | 15187 | Venezuela | F | *E. foetidum* |
| *E. foetidum* | V0471791F | D.M. White | 605 | Colombia | F | *E. foetidum* |
| *E. foetidum* | G.Davidse_15072 | G. Davidse | 15072 | Venezuela | K | *E. foetidum* |
| *E. foetidum* | T.Plowman_13505 | T. Plowman | 13505 | Venezuela | K | *E. foetidum* |
| *E. foetidum* | T.Plowman_13512 | T. Plowman | 13512 | Venezuela | K | *E. foetidum* |
| *E. foetidum* | P05576148 | E.P. Pinto | 1584 | Colombia | P | *E. foetidum* |
| *E. foetidum* | 2713682 | G. Davidse | 15072 | Venezuela | MO | *E. foetidum* |
| *E. foetidum* | VEN118524 | J. Steyermark | 113852 | Venezuela | VEN | *E. foetidum* |
| *E. foetidum* | VEN153973 | J. Steyermark | 122557 | Venezuela | VEN | *E. foetidum* |
| *E. gracilipes* | ASU0032976 | M. Baker | 7323 | Ecuador | ASU | *E. gracilipes* |
| *E. gracilipes* | 12621 | unreadable | 348 | Venezuela | B | *E. gracilipes* |
| *E. gracilipes* | BC624069 | J. Cuatrecasas | 15736 | Colombia | BC | *E. gracilipes* |
| *E. gracilipes* | BR0000005947157 | R. Spruce | 3068 | Venezuela | BR | *E. gracilipes* |
| *E. gracilipes* | BR000000699762 | R. Spruce | 3068 | Venezuela | BR | *E. gracilipes* |
| *E. gracilipes* | 627635 | G. Klug | 1440 | Peru | F | *E. gracilipes* |
| *E. gracilipes* | 690935 | G. Klug | 1961 | Colombia | F | *E. gracilipes* |
| *E. gracilipes* | 1193612 | L. Williams | 15561 | Venezuela | F | *E. gracilipes* |
| *E. gracilipes* | 1241355 | J. Cuatrecasas | 13303 | Colombia | F | *E. gracilipes* |
| *E. gracilipes* | 1365541 | J. Cuatrecasas | 15736 | Colombia | F | *E. gracilipes* |

| | | | | | | |
|---|---|---|---|---|---|---|
| *E. gracilipes* | 1756398 | T. Plowman | 2546 | Peru | F | *E. gracilipes* |
| *E. gracilipes* | 1757602 | T. Plowman | 2254 | Colombia | F | *E. gracilipes* |
| *E. gracilipes* | 1758849 | T. Croat | 18428 | Peru | F | *E. gracilipes* |
| *E. gracilipes* | 1774905 | T. Plowman | 2546 | Peru | F | *E. gracilipes* |
| *E. gracilipes* | 1842755 | I. Cabrera | 3637 | Colombia | F | *E. gracilipes* |
| *E. gracilipes* | 1845849 | J. Revilla | 2455 | Peru | F | *E. gracilipes* |
| *E. gracilipes* | 1850299 | L. Aristeguieta | 7111 | Venezuela | F | *E. gracilipes* |
| *E. gracilipes* | 1858321 | M.F. Silva | 38.841 | Brazil | F | *E. gracilipes* |
| *E. gracilipes* | 1867926 | M. Madison | 6471 | Brazil | F | *E. gracilipes* |
| *E. gracilipes* | 1873324 | M. Nee | 17507 | Venezuela | F | *E. gracilipes* |
| *E. gracilipes* | 1875165 | R. Liesner | 9210 | Venezuela | F | *E. gracilipes* |
| *E. gracilipes* | 1895285 | R. Liesner | 8977 | Venezuela | F | *E. gracilipes* |
| *E. gracilipes* | 1895311 | R. Liesner | 11530 | Venezuela | F | *E. gracilipes* |
| *E. gracilipes* | 1919610 | S. King | 430 | Peru | F | *E. gracilipes* |
| *E. gracilipes* | 1953854 | B. Stein | 2573 | Ecuador | F | *E. gracilipes* |
| *E. gracilipes* | 1959876 | W. Palacios | 348 | Ecuador | F | *E. gracilipes* |
| *E. gracilipes* | 1959877 | W. Palacios | 812 | Ecuador | F | *E. gracilipes* |
| *E. gracilipes* | 1962342 | R. Liesner | 17187 | Venezuela | F | *E. gracilipes* |
| *E. gracilipes* | 1964689 | G. Aymard | 4543 | Venezuela | F | *E. gracilipes* |
| *E. gracilipes* | 1965889 | L. Aristeguieta | 6353 | Venezuela | F | *E. gracilipes* |
| *E. gracilipes* | 1965890 | L. Williams | 15561 | Venezuela | F | *E. gracilipes* |
| *E. gracilipes* | 1975330 | B. Stergios | 5162 | Venezuela | F | *E. gracilipes* |
| *E. gracilipes* | 1982323 | B. Holst | 3146 | Venezuela | F | *E. gracilipes* |
| *E. gracilipes* | 1982366 | J. Brandbyge | 30871 | Ecuador | F | *E. gracilipes* |
| *E. gracilipes* | 1984265 | R. Vásquez | 2647 | Peru | F | *E. gracilipes* |
| *E. gracilipes* | 1984271 | R. Vásquez | 7142 | Peru | F | *E. gracilipes* |
| *E. gracilipes* | 1985367 | J. Brandbyge | 30816 | Ecuador | F | *E. gracilipes* |
| *E. gracilipes* | 1992274 | M. Baker | 6841 | Ecuador | F | *E. gracilipes* |
| *E. gracilipes* | 1993755 | C. Gerardo Aymard | 5368 | Venezuela | F | *E. gracilipes* |
| *E. gracilipes* | 2155567 | S. McDaniel | 29917 | Peru | F | *E. gracilipes* |
| *E. gracilipes* | 2198674 | J. Aronson | 888 | Peru | F | *E. gracilipes* |
| *E. gracilipes* | 2198675 | R. Vásquez | 9661 | Peru | F | *E. gracilipes* |
| *E. gracilipes* | 2198676 | R. Vásquez | 2040 | Peru | F | *E. gracilipes* |
| *E. gracilipes* | 2198678 | P.J.M. Maas | 6746 | Brazil | F | *E. gracilipes* |
| *E. gracilipes* | 2198679 | H.T. Beck | 427 | Brazil | F | *E. gracilipes* |
| *E. gracilipes* | 2211291 | M. Baker | 7323 | Ecuador | F | *E. gracilipes* |
| *E. gracilipes* | 2295279 | P. Acevedo-Rdgz. | 10348 | Venezuela | F | *E. gracilipes* |
| *E. gracilipes* | 2318236 | I. Huamantupa | 14277 | Peru | F | *E. gracilipes* |
| *E. gracilipes* | 2318474 | M. Ríos | 4353 | Peru | F | *E. gracilipes* |
| *E. gracilipes* | 2320258 | D.M. White | 476 | Peru | F | *E. gracilipes* |
| *E. gracilipes* | 2321133 | M. Ríos | 2696 | Peru | F | *E. gracilipes* |

| | | | | | | |
|---|---|---|---|---|---|---|
| *E. gracilipes* | 2324350 | D.M. White | 567 | Colombia | F | *E. gracilipes* |
| *E. gracilipes* | 2324355 | D.M. White | 566 | Colombia | F | *E. gracilipes* |
| *E. gracilipes* | 2324356 | D.M. White | 587 | Colombia | F | *E. gracilipes* |
| *E. gracilipes* | Bernardi_7459 | A.L. Bernardi | 7459 | Venezuela | K | *E. gracilipes* |
| *E. gracilipes* | K000073776 | H.C. Evans | FOX291 | Venezuela | K | *E. gracilipes* |
| *E. gracilipes* | K000424403 | R. Spruce | 3725 | Venezuela | K | *E. gracilipes* |
| *E. gracilipes* | K000424404 | R. Spruce | 3068 | Venezuela | K | *E. gracilipes* |
| *E. gracilipes* | K001201220 | D.B. Fanshawe | 6003 | Guyana | K | *E. gracilipes* |
| *E. gracilipes* | Kluge_1961 | G. Klug | 1961 | Colombia | K | *E. gracilipes* |
| *E. gracilipes* | s.n. | T.D.P. | 7626 | Ecuador | K | *E. gracilipes* |
| *E. gracilipes* | s.n. | B. Holst | 3146 | Venezuela | K | *E. gracilipes* |
| *E. gracilipes* | s.n. | C. Ellison | FOX243 | Ecuador | K | *E. gracilipes* |
| *E. gracilipes* | s.n. | J. Cuatrecasas | 13303 | Colombia | K | *E. gracilipes* |
| *E. gracilipes* | s.n. | J.G. Boer | 2345 | Venezuela | K | *E. gracilipes* |
| *E. gracilipes* | s.n. | M. Madison | 6471 | Brazil | K | *E. gracilipes* |
| *E. gracilipes* | s.n. | N. Jaramillo | 2647 | Peru | K | *E. gracilipes* |
| *E. gracilipes* | s.n. | R. Schultes | 6957 | Colombia | K | *E. gracilipes* |
| *E. gracilipes* | s.n. | W. Palacios | 2203 | Ecuador | K | *E. gracilipes* |
| *E. gracilipes* | s.n. | W. Palacios | 2573 | Ecuador | K | *E. gracilipes* |
| *E. gracilipes* | P00723632 | J. Cuatrecasas | 15736 | Colombia | P | *E. gracilipes* |
| *E. gracilipes* | P05483189 | F. Geay | 7197 | Venezuela | P | *E. gracilipes* |
| *E. gracilipes* | 6326091 | D.C. Daly | 8767 | Brazil | MO | *E. gracilipes* |
| *E. gracilipes* | s.n. | W. Palacios | 348 | Ecuador | MO | *E. gracilipes* |
| *E. gracilipes* | s.n. | R. Liesner | 21325 | Venezuela | MO | *E. gracilipes* |
| *E. gracilipes* | 1075681 | D.C. Daly | 11041 | Brazil | NY | *E. gracilipes* |
| *E. gracilipes* | 2171837 | P.J.M. Maas | 6670 | Brazil | NY | *E. gracilipes* |
| *E. gracilipes* | 2171839 | M. Nee | 34658 | Brazil | NY | *E. gracilipes* |
| *E. gracilipes* | 2171840 | L.S. Coelho | 10 | Brazil | NY | *E. gracilipes* |
| *E. gracilipes* | 2175394 | L.S. Coelho | 214 | Brazil | NY | *E. gracilipes* |
| *E. gracilipes* | 2817844 | A. Dik | 884 | Ecuador | NY | *E. gracilipes* |
| *E. gracilipes* | 2817863 | C.Aulestia | 30 | Ecuador | NY | *E. gracilipes* |
| *E. gracilipes* | 2864560 | K.M. Redden | 5357 | Guyana | NY | *E. gracilipes* |
| *E. gracilipes* | U0105998 | D.B. Fanshawe | 6003 | Guyana | U | *E. gracilipes* |
| *E. gracilipes* | U0106346 | A. Gentry | 41064 | Colombia | U | *E. gracilipes* |
| *E. gracilipes* | U0106347 | L. Aristeguieta | 7111 | Venezuela | U | *E. gracilipes* |
| *E. gracilipes* | 2772840 | J. Cuatrecasas | 15736 | Colombia | US | *E. gracilipes* |
| *E. gracilipes* | W0018333 | R. Spruce | 3068 | Venezuela | W | *E. gracilipes* |
| *E. gracilipes* | W18890163289 | R. Spruce | 3068 | Venezuela | W | *E. gracilipes* |
| *E. lineolatum* | s.n. | de Fremer | 1949 | Surinam | K | *E. lineolatum* |
| *E. lineolatum* | s.n. | Lehmann | s.n. | Colombia | K | *E. lineolatum* |
| *E. lineolatum* | s.n. | Maguire | 23337 | British Guiana | K | *E. lineolatum* |

| | | | | | | |
|---|---|---|---|---|---|---|
| *E. lineolatum* | s.n. | Burnham | 2714 | Surinam | MO | *E. lineolatum* |
| *E. n.* var. *novogranatense* | B100355177 | Schwerdtfeger | 7307 | Netherlands | B | *E. n.* var. *novogranatense* |
| *E. n.* var. *novogranatense* | COL000373550 | without collector | s.n. | Colombia | COL | *E. n.* var. *novogranatense* |
| *E. n.* var. *novogranatense* | COL000373551 | without collector | s.n. | Colombia | COL | *E. n.* var. *novogranatense* |
| *E. n.* var. *novogranatense* | G00352314 | A. Glaziou | 18160 | Brazil | G | *E. n.* var. *novogranatense* |
| *E. n.* var. *novogranatense* | GENT10167892 | J.J. Linden | 1181 | Colombia | GENT | *E. n.* var. *novogranatense* |
| *E. n.* var. *novogranatense* | K000424402 | Purdie | s.n. | Colombia | K | *E. n.* var. *novogranatense* |
| *E. n.* var. *novogranatense* | K000700849 | without collector | s.n. | UK | K | *E. n.* var. *novogranatense* |
| *E. n.* var. *novogranatense* | K000852894 | A. Glaziou | 18160 | Brazil | K | *E. n.* var. *novogranatense* |
| *E. n.* var. *novogranatense* | L2120931 | T.R. Chand | 5475 | India | L | *E. n.* var. *novogranatense* |
| *E. n.* var. *novogranatense* | L2120933 | J.C. Liao | 10699 | indet. | L | *E. n.* var. *novogranatense* |
| *E. n.* var. *novogranatense* | L2120935 | Idjan Nedi | 215 | Indonesia | L | *E. n.* var. *novogranatense* |
| *E. n.* var. *novogranatense* | L2120936 | S.M. Popta | 831/195 | Indonesia | L | *E. n.* var. *novogranatense* |
| *E. n.* var. *novogranatense* | L2120937 | S.M. Popta | 831/195 | Indonesia | L | *E. n.* var. *novogranatense* |
| *E. n.* var. *novogranatense* | L2120938 | W. Kaudern | 2 | Indonesia | L | *E. n.* var. *novogranatense* |
| *E. n.* var. *novogranatense* | L2120940 | unreadable | 8135 | Jamaica | L | *E. n.* var. *novogranatense* |
| *E. n.* var. *novogranatense* | L2120942 | without collector | s.n. | indet. | L | *E. n.* var. *novogranatense* |
| *E. n.* var. *novogranatense* | L2120950 | without collector | s.n. | indet. | L | *E. n.* var. *novogranatense* |
| *E. n.* var. *novogranatense* | L2120951 | V.F. Schiffner | 8 | indet. | L | *E. n.* var. *novogranatense* |
| *E. n.* var. *novogranatense* | L2120952 | without collector | s.n. | indet. | L | *E. n.* var. *novogranatense* |
| *E. n.* var. *novogranatense* | L3700971 | H.J. van Hattum | 995 | Colombia | L | *E. n.* var. *novogranatense* |
| *E. n.* var. *novogranatense* | L3783970 | H.H. Zeijlstra | s.n. | indet. | L | *E. n.* var. *novogranatense* |
| *E. n.* var. *novogranatense* | MA-01-00459514 | T. Plowman | 6179 | USA | MA | *E. n.* var. *novogranatense* |
| *E. n.* var. *novogranatense* | MA-01-00459515 | T. Plowman | 7626 | USA | MA | *E. n.* var. *novogranatense* |
| *E. n.* var. *novogranatense* | MA-01-00663401 | without collector | s.n. | Colombia | MA | *E. coca* var. *coca* |
| *E. n.* var. *novogranatense* | MA-01-00663402 | without collector | s.n. | Colombia | MA | *E. n.* var. *novogranatense* |
| *E. n.* var. *novogranatense* | MA-01-00663403 | without collector | s.n. | Colombia | MA | *E. n.* var. *novogranatense* |
| *E. n.* var. *novogranatense* | MA-01-00663404 | without collector | s.n. | Colombia | MA | *E. n.* var. *novogranatense* |
| *E. n.* var. *novogranatense* | MA-01-00663405 | without collector | s.n. | Colombia | MA | *E. n.* var. *novogranatense* |

| | | | | | | |
|---|---|---|---|---|---|---|
| *E. n.* var. *novogranatense* | MA-01-00895854 | J. Cuatrecasas | 24310 | Colombia | MA | *E. n.* var. *novogranatense* |
| *E. n.* var. *novogranatense* | MA-02-00255978 | T. Plowman | 10980 | Peru (grown from seeds collected in Colombia) | MA | *E. n.* var. *novogranatense* |
| *E. n.* var. *novogranatense* | MA663401 | without collector | s.n. | Colombia | MA | *E. coca* var. *coca* |
| *E. n.* var. *novogranatense* | P00723942 | M.A. Glaziou | 18160 | Brazil | P | *E. n.* var. *novogranatense* |
| *E. n.* var. *novogranatense* | P05482996 | T. Plowman | 7626 | USA | P | *E. n.* var. *novogranatense* |
| *E. n.* var. *novogranatense* | P05482999 | J. Goudot | s.n. | Colombia | P | *E. n.* var. *novogranatense* |
| *E. n.* var. *novogranatense* | P05483000 | J. Triana | s.n. | Colombia | P | *E. n.* var. *novogranatense* |
| *E. n.* var. *novogranatense* | P05483007 | J.H. Hart | 5830 | Trinidad and Tobago | P | *E. n.* var. *novogranatense* |
| *E. n.* var. *novogranatense* | P05483012 | T. Plowman | 6199 | USA | P | *E. n.* var. *novogranatense* |
| *E. n.* var. *novogranatense* | P05483013 | T. Plowman | 8035 | USA | P | *E. n.* var. *novogranatense* |
| *E. n.* var. *novogranatense* | P05483014 | without collector | s.n. | France | P | *E. n.* var. *novogranatense* |
| *E. n.* var. *novogranatense* | NCSC00014120 | E. Machado | 2997 | Peru | NCSC | *E. n.* var. *novogranatense* |
| *E. n.* var. *novogranatense* | U0001831 | without collector | s.n. | indet. | U | *E. n.* var. *novogranatense* |
| *E. n.* var. *novogranatense* | U1284624 | J.D. van Leeuwen-Reijnvaan | s.n. | indet. | U | *E. n.* var. *novogranatense* |
| *E. n.* var. *novogranatense* | U1284625 | J. Heylen | s.n. | Belgium | U | *E. n.* var. *novogranatense* |
| *E. n.* var. *novogranatense* | U1284628 | without collector | s.n. | indet. | U | *E. n.* var. *novogranatense* |
| *E. n.* var. *novogranatense* | U1284629 | M. Buysman | 7 | Grenada | U | *E. n.* var. *novogranatense* |
| *E. n.* var. *novogranatense* | U1284665 | T. Plowman | 10980 | Peru | U | *E. n.* var. *novogranatense* |
| *E. n.* var. *novogranatense* | U1284666 | T. Plowman | 3768 | Colombia | U | *E. n.* var. *novogranatense* |
| *E. n.* var. *novogranatense* | U1284668 | P.J.M. Maas | 2036 | Colombia | U | *E. n.* var. *novogranatense* |
| *E. n.* var. *novogranatense* | U1284670 | T. Plowman | 7670 | Venezuela | U | *E. coca* var. *coca* |
| *E. n.* var. *novogranatense* | U1284672 | P.J.M. Maas | 1886 | Colombia | U | *E. n.* var. *novogranatense* |
| *E. n.* var. *novogranatense* | U1284673 | P.J.M. Maas | 1836 | Colombia | U | *E. coca* var. *coca* |
| *E. n.* var. *novogranatense* | WAG1930538 | J.J. Bos | 1488 | Netherlands | WAG | *E. n.* var. *novogranatense* |
| *E. n.* var. *novogranatense* | WAG1930539 | J.J. Bos | 1488 | Netherlands | WAG | *E. n.* var. *novogranatense* |
| *E. n.* var. *novogranatense* | WAG1930540 | G.H. Ruisch | 8484 | Netherlands | WAG | *E. n.* var. *novogranatense* |
| *E. n.* var. *novogranatense* | WAG1930542 | J.J. de Wilde | 12 | USA | WAG | *E. n.* var. *novogranatense* |
| *E. n.* var. *novogranatense* | WAG1930552 | T. Plowman | 1096 | USA | WAG | *E. n.* var. *novogranatense* |

| | | | | | | |
|---|---|---|---|---|---|---|
| *E. n.* var. *truxillense* | 168065 | T. Plowman | 6239 | USA | FLAS | *E. n.* var. *truxillense* |
| *E. n.* var. *truxillense* | L2120954 | T. Plowman | 6243 | USA | L | *E. n.* var. *truxillense* |
| *E. n.* var. *truxillense* | MA-01-00459516 | T. Plowman | 6239 | USA | MA | *E. n.* var. *truxillense* |
| *E. n.* var. *truxillense* | MA-01-00459517 | T. Plowman | 6234 | USA | MA | *E. n.* var. *truxillense* |
| *E. n.* var. *truxillense* | P05483003 | M.T Madison | 4920 | Ecuador | P | *E. n.* var. *truxillense* |
| *E. n.* var. *truxillense* | P05483004 | T. Plowman | 5607 | Peru | P | *E. n.* var. *truxillense* |
| *E. n.* var. *truxillense* | P05483005 | T. Plowman | 6204 | USA | P | *E. n.* var. *truxillense* |
| *E. n.* var. *truxillense* | P05483006 | T. Plowman | 6225 | USA | P | *E. n.* var. *truxillense* |
| *E. n.* var. *truxillense* | NCSC00014121 | E. Machado | 2573 | Peru | NCSC | *E. n.* var. *truxillense* |
| *E. n.* var. *truxillense* | NCSC00014122 | E. Machado | 1214 | Peru | NCSC | *E. n.* var. *truxillense* |
| *E. n.* var. *truxillense* | NCSC00014123 | E. Machado | 1239 | Peru | NCSC | *E. n.* var. *truxillense* |
| *E. n.* var. *truxillense* | NCSC00014124 | E. Machado | 1051 | Peru | NCSC | *E. n.* var. *truxillense* |
| *E. n.* var. *truxillense* | NCSC00014125 | E. Machado | 1201 | Peru | NCSC | *E. n.* var. *truxillense* |
| *E. n.* var. *truxillense* | NCSC00014126 | E. Machado | 2963 | Peru | NCSC | *E. n.* var. *truxillense* |
| *E. n.* var. *truxillense* | NCSC00014127 | E. Machado | 2960 | Peru | NCSC | *E. n.* var. *truxillense* |
| *E. n.* var. *truxillense* | NCSC00014128 | E. Machado | 2987 | Peru | NCSC | *E. n.* var. *truxillense* |
| *E. n.* var. *truxillense* | NCSC00014129 | E. Machado | 1018 | Peru | NCSC | *E. n.* var. *truxillense* |
| *E. n.* var. *truxillense* | NCSC00014130 | E. Machado | 1248 | Peru | NCSC | *E. n.* var. *truxillense* |
| *E. n.* var. *truxillense* | U1284634 | T. Plowman | 6209 | USA | U | *E. n.* var. *truxillense* |
| *E. n.* var. *truxillense* | UTC00203669 | T. Plowman | 13831 | USA | UTC | *E. n.* var. *truxillense* |
| *E. n.* var. *truxillense* | WAG1930553 | T. Plowman | 6243 | USA | WAG | *E. n.* var. *truxillense* |
| *E. n.* var. *truxillense* | WAG1930554 | T. Plowman | 6205 | USA | WAG | *E. n.* var. *truxillense* |

**Table S2. Taxon information for all specimens sequenced for this study.**

| Taxon | Year of Collection | Collector | Collector Number | Country of Collection | Type? | K Accession |
|---|---|---|---|---|---|---|
| *E. gracilipes* | 1853 | R. Spruce | 3068 | Venezuela | Isotype | K000424404 |
| *E. gracilipes* | 1949 | T. Landing | 6003 | British Guiana | | K001201220 |
| *E. gracilipes* | 1994 | H. C. Evans, D. R. Varlev | 291 | Venezuela | | K000073776 |
| *E. gracilipes* | 1987 | B.K. Holst & R. Liesner | 3146 | Venezuela | | |
| *E. gracilipes* | 1987 | R. Liesner & B.K. Holst | 21325 | Venezuela | | |

| | | | | | | |
|---|---|---|---|---|---|---|
| *E. gracilipes* | 1981 | R. Vásquez & N. Jaramillo | 2647 | Peru | | |
| *E. gracilipes* | 1993 | C.A. Ellison, & H.C. Evans | 243 | Ecuador | | |
| *E. gracilipes* | 1985 | J. Zaruma, W. Palacios | 2573 | Ecuador | | |
| *E. gracilipes* | 1931 | G. Klug | 1961 | Colombia | | |
| *E. gracilipes* | 1962 | C.D. Cazalet & T.D. Pennington | 7626 | Ecuador | | |
| *E. gracilipes* | 1941 | J. Cuatrecasas | 13303 | Colombia | | |
| *E. gracilipes* | 1985 | W. Palacios | 348 | Ecuador | | |
| *E. gracilipes* | 1853? | R. Spruce | 3725 | Venezuela | Lectotype | K000424403 |
| *E. cataractarum* | 1854 | R. Spruce | 3585 | Venezuela | | K001201221 |
| *E. cataractarum* | 1854 | R. Spruce | 3565 | Venezuela | | K001201222 |
| *E. cataractarum* | 1852 | R. Spruce | 2614 | Brazil | Holotype | K000426845 |
| *E. cataractarum* | 1941 | J. Cuatrecasas | 13277 | Colombia | | |
| *E. lineolatum* | 1906 | s.n. | 4736 | Colombia | | |
| *E. gracilipes* | 1978 | M.T. Madison,, A.H Kennedy, O. Monteiro, P.I.S. Braga | 6471 | Brazil | | K001201219 |
| *E. lineolatum* | 1944 | B. Maguire, D.B. Fanshawe, | 23337 | British Guiana | | |
| *E. foetidum* | 1984 | T. Plowman, F. Guánchez | 13512 | Venezuela | | |
| *E. novogranatense* | 2019 | S. S. Renner | 2888 | Munich Botanical Garden, Germany | | |

**Table S3. SRR accession numbers and taxon information for specimens used in the plastid phylogeny.**

| SRR accession number | Taxon | Year of Collection | Collector | Collector Number | Country of Collection | Accession |
|---|---|---|---|---|---|---|
| SRR7749349 | *E. novogranatense* | 1981 | T. Plowman | 10980 | Peru | *F_1898296* |
| SRR7749357 | *E. gracilipes* | 1987 | H. T. Beck, B. Rabelo., R. Souza, D. Seriacu. | 427 | Brazil | *F_2198679* |
| SRR7749361 | *E. gracilipes* | 1987 | B. K. Holst& R. Liesner | 3146 | Venezuela | *F_1982323* |
| SRR7749397 | *E. novogranatense* var. *truxillense* | | T. Plowman | 5600 | Peru | *F_1873746* |
| SRR7749404 | *E. cataractarum* | 1987 | R. Callejas | 471 | Colombia | *F_2198726* |
| SRR7749407 | *E. coca ipadu* | 1980 | T. Plowman | 6923 | Peru | *F_1823932* |
| SRR7749409 | *E. coca coca* | 1988 | M. Lewis | 36915 | Bolivia | *F_2138572* |
| SRR7749410 | *E. coca coca* | 2013 | - | - | USA | *USDA_B145* |
| SRR7749418 | *E. novogranatense* | 1978 | T.C. | 7670 | Venezuela | *F_1853454* |

**Table S4. Degree of correspondence between taxon-designated groups and GMM-designated groups, linear metric datasets.**

| **Linear metric dataset** | **Rand index** |
|---|---|
| main dataset (835 leaves) | 0.691 |
| cultivated only (404 leaves) | 0.632 |
| cultivated, one leaf per specimen (97 leaves) | 0.595 |

**Table S5. Degree of correspondence between taxon-designated groups and GMM-designated groups, EFA datasets.**

| **EFA dataset** | **Rand index** |
|---|---|
| full dataset (835 leaves) | 0.555 |
| cultivated only (404 leaves) | 0.425 |
| cultivated, one leaf per specimen (97 leaves) | 0.450 |

### Supplementary Results

##### *Morphometric analysis*

Although recent advances in computation and statistics have allowed researchers to apply quantitative methods to the study of shape, superseding qualitative descriptions in their precision (Jensen, 2003; Claude, 2008), no such approach has been attempted for coca and its relatives. Collectively, our morphometric results show that of all wild relatives studied, *E. cataractarum* is most similar in its leaf size and shape to *E. coca* and *E. novogranatense.* Therefore, morphological discrimination of wild from cultivated varieties is also not reliable using EFA (shape) metrics. The homogeneity of *E. n.* var. *truxillense* groups provides tentative support for this being a variety with either a narrower gene pool or a lower phenotypical plastic response, occupying a narrow portion of the morphospace. Overall, there is less uniformity of taxa when grouped using shape data compared to linear metric data This implies considerable phenotypic plasticity in leaf shape driven by variation of environmental, stochastic and interacting genetic effects (Chitwood & Sinha, 2016).

##### *Phylogenomic analysis*

The earliest diverging clade on the respective nuclear and plastid trees both contain the *E. gracilipes* Spruce 3725 lectotype from Venezuela and *E. foetidum* as a sister taxon regardless of which sample of *E. foetidum* was used. In the nuclear tree, the *E. gracilipes* 3068 isotype appears in the *E. coca* + *E. gracilipes* clade, as sister to sample *E. gracilipes* 6471 from Brazil (86% quartet support). *E. gracilipes* samples placed differently in the plastid compared to the nuclear tree suggest gene flow involving this species. In the plastid tree, the *E. gracilipes* 3068 isotype forms the earliest-diverging branch in the *E. novogranatense* clade, while in the nuclear tree, this isotype is placed in the *E. coca* + *E. gracilipes* clade. Further exemplifying the incongruence are the two geographical clades of *E. gracilipes* (Ecuador, *n*=3 and Venezuela, *n*=2). Forming a monophyletic group in the plastid tree with 100% bootstrap support, these are separated in the ASTRAL nuclear summary tree, in which the Ecuador group is the earliest diverging (after *E. foetidum*). This bipartition has more supporting trees (92) than conflicting trees (69) and moderate gene tree support values (gCF= 0.57, IC=0.42). The high level of overall discordance implies that the *E. coca* + *E. gracilipes* clade constitutes a recent radiation with incomplete lineage sorting.

Two samples could not be compared due to a low retrieval success of nuclear genes and their consequent omittance from the nuclear tree (*E. gracilipes* 13303 – 61 genes and *E. gracilipes* 21325 – 28 genes). When these two samples were added to the plastid dataset, they fell within the Ecuador/ Venezuela clade, with *E. gracilipes* 13303 (Colombia) as the earliest diverging branch (Fig. S11).

##### *Lineage dating analysis*

StarBEAST2 also produced an alternative species tree topology, albeit with much reduced support. In this topology, *E. novogranatense* is placed as sister to the remainder of the coca clade (1.0 PP), and *E. cataractarum* as sister to *E. coca* var. *coca* and *E. gracilipes* (0.3 PP) (Fig. 5).
